# A precise and adaptive neural mechanism for predictive temporal processing in the frontal cortex

**DOI:** 10.1101/2021.03.10.434831

**Authors:** Nicolas Meirhaeghe, Hansem Sohn, Mehrdad Jazayeri

## Abstract

The theory of predictive processing posits that the nervous system uses expectations to process information predictively. Direct empirical evidence in support of this theory however has been scarce and largely limited to sensory areas. Here, we report a precise and adaptive neural mechanism in the frontal cortex of non-human primates consistent with predictive processing of temporal events. We found that the speed at which neural states evolve over time is inversely proportional to the statistical mean of the temporal distribution of an expected stimulus. This lawful relationship was evident across multiple experiments and held true during learning: when temporal statistics underwent covert changes, neural responses underwent predictable changes that reflected the new mean. Together, these results highlight a precise mathematical relationship between temporal statistics in the environment and neural activity in the frontal cortex that could serve as a mechanistic foundation for predictive temporal processing.

## Introduction

Since the early work of Hermann Helmholtz (*1*), the field of psychology has embraced the idea that humans rely on prior knowledge to construct a coherent interpretation of raw sensory inputs. Central to this idea is the notion of internal models that represent statistical regularities in the environment (*2–4*) and help us make better perceptual inferences (*5*), optimize behavioral responses (*6*), and swiftly adapt to environmental changes (*7*).

Internal models establish lawful relationships between statistical regularities in the environment and neural signals in the brain (*8–10*). However, the inherent complexities of circuits and signals in the nervous system have made the discovery and concrete characterization of such lawful relationships extremely challenging (*11, 12*). One influential hypothesis inspired by information-theoretic accounts of brain function (*13, 14*) is that prior statistics enable the brain to process incoming information predictively; i.e., in terms of deviations from expectations (*15–17*). To do so, the nervous system is thought to encode the predictable components of sensory inputs, which are compared to the actual inputs once they become available (*18, 19*). Predictive processing has been touted as a canonical cortical computation and used to provide a functional account of a wide range of cortical phenomena, including the response properties of neurons in early visual cortex (*20*) and the logic of laminar information processing in cortical circuits (*21, 22*).

The conceptual impact of this theory however has been far greater than evidence in its support. Here, we highlight three major gaps in our current understanding that the present study aims to address. First, high-resolution recordings in animal experiments have largely focused on how individual neurons in sensory areas encode sensory predictions (*11, 16, 23, 24*). For example, the foundational implications of predictive processing for efficient coding (*13*) have only been explored in terms of tuning properties of spatially-tuned sensory neurons (*25*). As such, evidence for predictive processing in higher-order cortical areas is still wanting (*26*). Second, the implications of predictive processing have typically been examined in terms of static firing rates of neurons and not their dynamics. Recent advances, in contrast, suggest that fundamental neural computations might be carried out through dynamic patterns of activity that emerge from interactions among populations of neurons (*27–34*). Therefore, it is critical that we revisit predictive internal models in terms of population coding and dynamics. Finally, one strong appeal of the theory is that it makes quantitative predictions about the relationship between neural signals and environmental statistics. Experimentally however, it has proven challenging to establish such precise relationships, even for simple statistical properties such as the expected value of a stochastic variable.

Here, we begin to address these outstanding questions based on analyses of neural activity in the frontal cortex of non-human primates. Using both old and new data, we uncover a precise mathematical relationship between neural activity and environmental statistics in the temporal domain: the speed at which neural responses evolve over time is inversely proportional to the average time that a stimulus is expected to occur. This lawful relationship was present across multiple independent experiments including a novel learning experiment wherein neural responses underwent predictable adjustments in accordance with changes in the mean of the distribution.

## Results

### Behavioral and neural signatures of predictive processing in the ‘Ready-Set-Go’ task

In a previous study, we trained two monkeys to perform a time interval reproduction task known as ‘Ready-Set-Go’ (RSG) (*28*) (Figure 1A). This task requires animals to (1) fixate a central spot, (2) measure a sample time interval (*t_s_*) between two visual flashes (‘Ready’ followed by ‘Set’), and (3) produce a matching interval (*t_p_*) immediately after Set by initiating a saccade (‘Go’) toward a visual target presented left or right of the fixation point. Animals performed this task in two conditions associated with two distinct *t_s_* distributions (Figure 1A**, top right**). The conditions were cued explicitly by the color of the fixation spot and were interleaved across short blocks of trials (Methods). When the fixation spot was red, *t_s_* was sampled from a ‘Short’ uniform distribution (480–800 ms). When the fixation spot was blue, *t_s_* was sampled from a ‘Long’ distribution (800–1200 ms). The amount of reward monkeys received at the end of each trial decreased linearly with the magnitude of the relative error ((*t_p_-t_s_*)/*t_s_*) (Figure 1A**, bottom left**).

**Figure 1.**
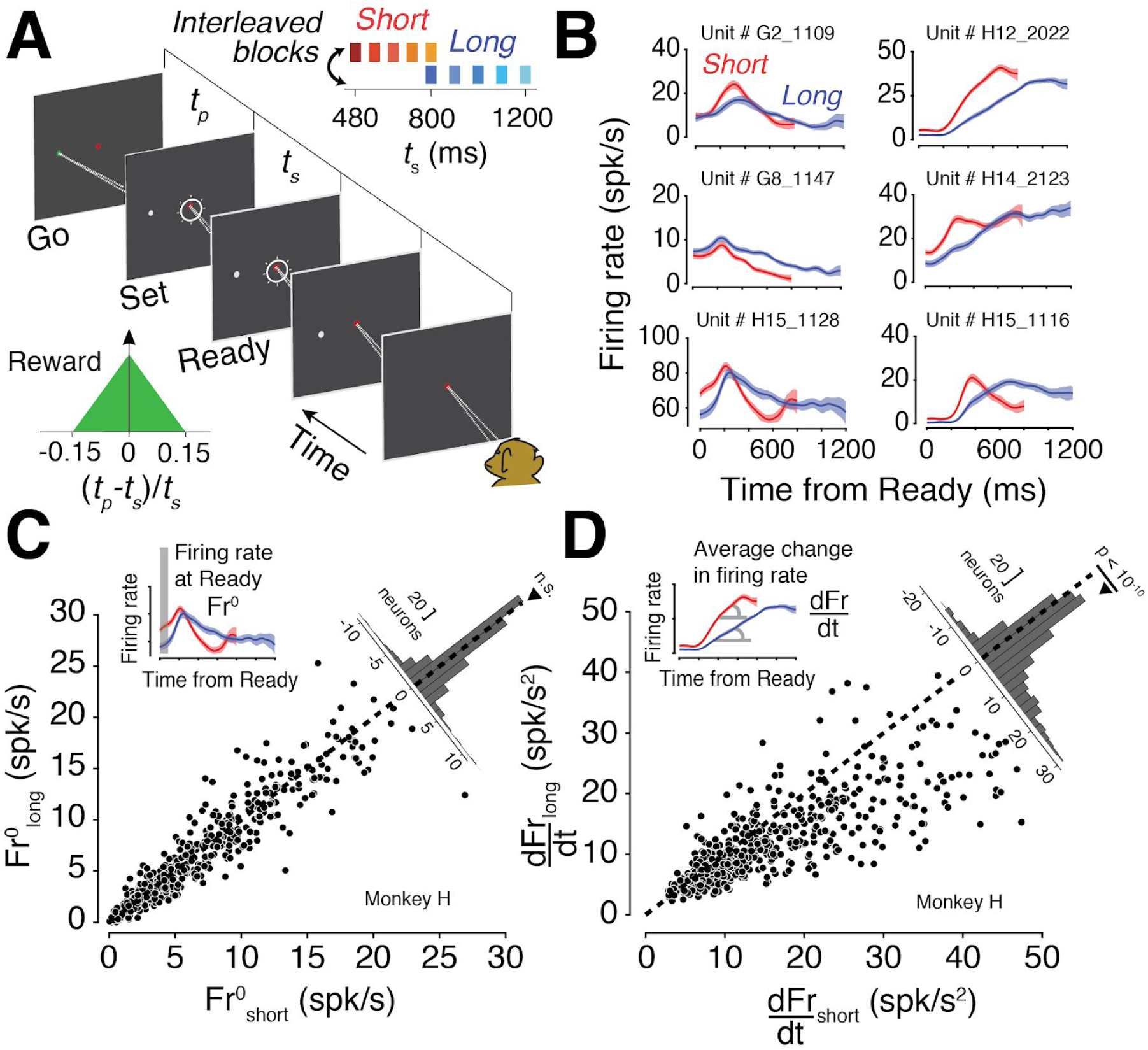
Single neuron signatures of temporal expectations during time estimation. **(A)** Ready-Set-Go task. On every trial, the monkey measures a sample interval (*t_s_*) between the Ready and Set cue (visual annulus flashed around the fixation point). After Set, the animal produces a matching interval (*t_p_*) by making a delayed saccade to a peripheral target (Go). Bottom left: reward function. The animal receives reward when *t_p_* is within a fixed window around *t_s_*. Inside this window, the amount of reward decreases linearly with the relative error (*t_p_*-*t_s_*)/*t_s_*. Top right: sample interval distributions. To modulate animals’ temporal expectations about the timing of the Set cue, *t_s_* was sampled from one of two distributions Short (red) or Long (blue). The distributions were interleaved in short blocks of trials (length: 4.0 ± 4.4 trials; uniform hazard) and indicated on every trial by the color of the fixation point. **(B)** Firing rate of six example neurons during the measurement epoch of the task, color-coded by condition. Shaded areas denote 95% CI obtained from bootstrap resampling (N=100). Firing rates were obtained after binning (*w_bin_*=20ms) and smoothing (*sd_kernel_*=40ms) spike counts averaged across trials. **(C)** Firing rate at the time of Ready (*Fr^0^*) for the Long versus Short condition. Each circle represents one neuron (N=619 for monkey G, N=542 for monkey H). *Fr^0^* was computed in a 20-ms window immediately following Ready. The diagonal distribution shows the difference (*Fr^0^_short_* - *Fr^0^_long_*) across the population of recorded neurons (paired t-test, *p*=0.40 for monkey G, *p*=0.07 for monkey H). **(D)** Change in instantaneous firing rate (*dFr/dt*) averaged within the measurement epoch for the Long versus Short condition. Each circle represents one neuron. *dFr/dt* was computed as the absolute difference in firing rate between consecutive 20-ms bins average over the measurement epoch, normalized by the bin size. Results were qualitatively unchanged if the duration of the averaging period in the Long condition was matched to that in the Short condition (i.e., averaging from Ready to Ready+800ms in both conditions). The diagonal distribution shows that the change in firing rate (i.e., ‘speed’) across the population of neurons is significantly higher in the Short compared to the Long condition (paired t-test, *p*<10^-10^ for both monkeys).

A salient feature of behavior in this task is that responses regress toward the mean of the interval distribution (*35*). We found a robust expression of this regression-to-the-mean phenomenon (*28*) causing opposite biases in the Short and Long conditions for the overlapping 800 ms interval (Figure S1). This observation suggests that animals measure time intervals predictively, i.e., relative to the mean of the temporal distribution. Accordingly, we hypothesized that the neural activity in the measurement epoch of RSG (i.e., between Ready and Set) encodes the mean of the interval distribution.

To test this hypothesis, we analyzed neural activity (N=619 neurons for monkey G, N=542 for monkey H) in the dorsomedial region of the frontal cortex (DMFC; Figure S2), an area that has been strongly implicated in providing temporal control over behavior in humans (*36–41*), monkeys (*42–50*), and rodents (*51–56*). During the measurement epoch, neurons had complex and heterogeneous response profiles that typically differed between the two conditions (Figure 1B, S3). Previous recordings in sensory and sensorimotor areas have reported a signature of prior expectations in terms of systematic changes in the baseline firing rate of individual neurons(*57–63*). Inspired by these previous findings, we performed a variety of analyses looking for systematic firing rate differences between the two conditions. At the level of individual neurons, differences in firing rates changed over time, and were unstructured (Figure S4) with activity sometimes stronger in the Short condition, and other times in the Long condition. Similarly, the normalized difference in firing rates across the population did not systematically differ between the two conditions (Figure S4). We also analyzed the difference between firing rates at the time of Ready, denoted Fr^0^, when the only information available to the animal was the condition type. Although many neurons had different levels of activity depending on the condition, across the population, neurons were equally likely to be more or less active in the Short compared to the Long condition (two-sided paired-sample t-test, *t*(617)=0.83, *p*=0.40 for monkey G, *t*(553)=-1.84, *p*=0.07 for monkey H, Figure 1C). Based on these results, we concluded that the expectation of the mean is not encoded by systematic firing rate differences across the two conditions.

Moving beyond static firing rates, we considered the possibility that the mean interval might be encoded in how firing rates change over time (i.e., neural dynamics). Recent work has highlighted the importance of neural dynamics in a wide array of tasks (*29, 32, 33, 64*). For example, it has been shown that neural responses compress or stretch in time according to previously learned temporal contingencies (*47, 48, 65, 66*), a phenomenon referred to as *temporal scaling*. Scaling has been reported during the planning of delayed movements (*51, 55, 67–70*), but also when animals anticipate an external event (*54, 71–73*). Neural responses in RSG exhibited a qualitatively similar temporal scaling between the two conditions; i.e., the activity profile in the Long condition often appeared as a stretched version of the activity profile in the Short condition (Figure 1B). To quantify this scaling effect, we estimated the average rate of change (i.e., ‘speed’) of firing rate over the measurement epoch for each neuron (Methods). Consistent with our qualitative observation, the speed across the population of neurons was on average larger in the Short compared to the Long condition (two-sided paired-sample t-test, *t*(617)=12.6, *p*<10^-10^ for monkey G, *t*(545)=12.9, *p*<10^-10^ for monkey H; Figure 1D, S4). This finding suggests that although prior expectations were not encoded by an overall modulation of firing rates, they impacted the speed at which neural responses evolved over time.

### Temporal scaling lawfully reflects the mean of experienced temporal distributions

Although the phenomenon of temporal scaling is not new (*47, 48, 51, 54, 55, 65–72, 74, 75*), its functional implication is not understood. Different studies have offered an interpretation in terms of attention (*76*), anticipation (*77, 78*), or reward expectation (*71, 79*). These interpretations however are largely qualitative and do not specify the exact relationship between temporal scaling and experimentally-imposed temporal statistics.

Here, we explore an alternative functional interpretation of temporal scaling that is grounded in the theory of predictive processing. According to this theory, neural signals ought to have a precise relationship to the statistics of sensory inputs: neural signals should represent what is expected (*11, 20, 25*) so that incoming information can be encoded relative to that expectation (*15–17*). Accordingly, we hypothesized that temporal scaling in the Ready-Set epoch is adjusted in proportion to the mean of the interval distribution. We refer to this hypothesis as the mean-predictive-temporal-scaling (MPTS) hypothesis. MPTS makes a specific prediction: the speed at which neural activity evolves over time must be inversely proportional to the average time that the stimulus is expected to occur (faster for earlier, slower for later). Equivalently, the ratio of speeds across different interval distributions should be exactly equal to the reciprocal ratio of the corresponding interval means.

We tested this prediction first at the level of single neurons. If patterns of activity are scaled according to the average interval, one should be able to reconstruct the firing rate of a neuron in the Short condition based on its activity in the Long condition by applying a precise scaling operation (Figure 2A). Let us call *r_short_*(*t*) and *r_long_*(*t*) the firing rates of a neuron in the Short and Long conditions, and μ*_short_* and μ*_long_*, the corresponding mean intervals. Our hypothesis predicts that r*_short_*(*t*) should be well approximated by *r_long_* (λ*t*), where the scaling factor λ is equal to the ratio 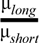, which in our experiment, is 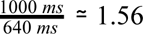. We thus fitted each neuron’s activity profile in the Short condition with a scaled version of its activity profile in the Long condition (see Methods and Figure S3, S6), and analyzed the fit to λ across the population.

**Figure 2.**
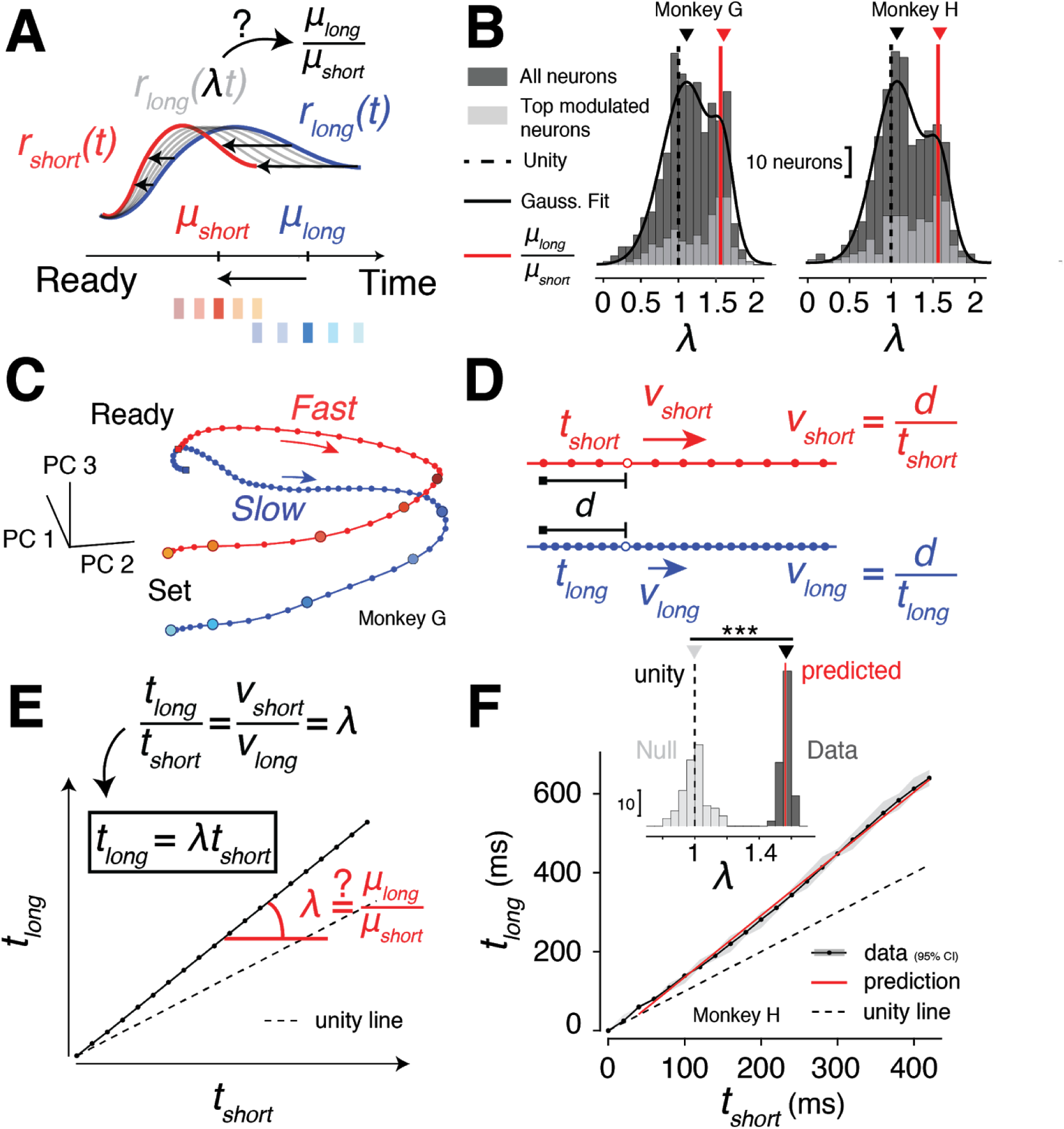
Neural dynamics encodes the statistical mean of the distribution. **(A)** Schematic demonstration of the temporal scaling analysis for individual neurons. We fitted each neuron’s response profile in the Short condition (*r_short_*(*t*), red) by a temporally-scaled version of the response in the Long condition (*r_long_* (*t*), blue). We predicted that the optimal scaling factor, λ, of the fitted response (*r_long_* (λ*t*), grey) would be equal to the ratio of the means of the two distributions, μ*_long_* /μ*_short_*. **(B)** Distribution of scaling factors across neurons in both monkeys (black: all neurons; grey: only neurons whose responses accounted for more than 0.1% of the total variance). Both the black and gray distributions had two peaks, one near unity (black dashed line) and one near the predicted value μ*_long_* /μ*_short_* (red line). We fitted a Gaussian mixture model to the black distribution (black line) to estimate the location of the two peaks (Methods). The first peak (M1, black triangle) was near unity (1.11±0.33 for monkey G, and 1.07±0.27 for monkey H; mean±sd), and the second peak (M2, red triangle) was near the predicted value (1.60±0.14 for monkey G, and 1.58±0.15 for monkey H). **(C)** Population dynamics during the measurement epoch. We applied principal component analysis (PCA) to visualize neural trajectories associated with both conditions in the subspace spanned by the top 3 PCs (∼75% of total variance explained). Population dynamics described two parallel trajectories evolving at different speeds, as shown qualitatively by the spacing between consecutive states (bin size: 20 ms) along the trajectories (faster for Short, in red, compared to Long, in blue). **(D)** Schematic demonstration of the temporal scaling analysis across the population. We estimated speed differences between the two conditions by computing the time needed to travel the same arbitrary distance *d* (black segment) along the Short and Long trajectory (*t_short_* and *t_long_*, respectively), starting from a common reference state (black square). Since immediately after Ready, responses were dominated by a non-specific, likely visually-evoked transient (Figure S4), we chose 400 ms after Ready as the reference state. However, the results were robust to the choice of reference state (Figure S5). **(E)** For any distance *d* traveled along the trajectories, we have by definition *v_short_* = *d/t_short_* and *v_long_* = *d*/*t_long_*. The ratio of speeds must therefore verify the relationship *v_short_*/*v_long_* = *t_long_* /*t_short_*. If the ratio of speeds also abides by *v_short_*/*v_long_* = μ_*long*_ /μ_*short*_ predicted by the MPTS hypothesis, then it follows that *t_long_* /*t_short_* = μ*_long_* /μ*_short_*, or equivalently, *t_long_* = (μ*_long_* /μ*_short_*) *t_short_*. Therefore, MPTS predicts that the mapping between *t_long_* and *t_short_* should be linear, and the slope of this mapping should be precisely equal to the predicted scaling factor, λ = μ*_long_* /μ*_short_*. **(F)** We plotted the mapping between *t_long_* and *t_short_* to estimate the empirical scaling factor relating the two neural trajectories. The mapping was linear and diverged from the unity line with a slope close to the predicted value μ*_long_* /μ*_short_* ≃ 1.56 (95% CI for scaling factor: [1.39 1.58] for monkey G, [1.50 1.64] for monkey H). Overlaid with the mapping, the red line shows the prediction constrained to have a slope equal to the value μ*_long_* /μ*_short_*; the intercept is chosen for visualization to minimize the root-mean-squared-error between the prediction and the data. Inset: the slope of a regression model on the empirical mapping (black distribution for bootstrapped values) matches the predicted value (red line) and differs significantly from the null value obtained by randomly shuffling conditions separately for each neuron (grey distribution).

We found that the distribution of scaling factors across the population was bimodal, with one peak near λ = 1 (no scaling), and another peak near the value predicted by MPTS, 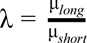 (Gaussian mixture model, mean±sd, M_1_=1.11±0.33, M_2_=1.60±0.14 for monkey G; M_1_=1.07±0.27, M_2_=1.58±0.15 for monkey H; Figure 2B). This bimodality suggested that the population comprised a mixture of two ensembles, a non-scaling and a scaling ensemble. To ensure that this dichotomy was present across task-modulated neurons, we repeated the same analysis across the subset of neurons which had the largest firing rate modulations during the measurement epoch. We sorted neurons by the amount of variance in their firing rate across time points in the measurement epoch and kept neurons which contributed more than 0.1% of the total variance (150/741 neurons for monkey G, 164/617 for monkey H; see Methods). The resulting distribution of scaling factors for this reduced population showed the same bimodality (M_1_=1.03±0.41, M_2_=1.58±0.18 for monkey G; M_1_=1.17±0.36, M_2_=1.59±0.16 for monkey H; Figure 2B). Together, these results provided evidence that the activity profile of a subpopulation of neurons in DMFC was exactly scaled according to the mean of the interval distribution.

Because the amount of scaling varied from neuron to neuron, we next sought to quantify the scaling effect at the population level. Following recent practices for high-dimensional neural datasets, we considered neural activity across the entire population as a state evolving in a high-dimensional state space where each dimension represents the activity of one neuron (*64, 80*). We first applied principal component analysis (PCA) to visualize how the dynamics in the two conditions unfolded between Ready and Set (Methods). Qualitatively, results were consistent with MPTS: the speed of neural trajectories appeared faster for the Short compared to the Long condition (Figure 2C).

To test whether the speeds were quantitatively consistent with MPTS, we devised an analysis to measure speed differences between conditions (Figure 2D; see Methods). Briefly, we computed the time (*t_short_* and *t_long_*) necessary to travel the same distance along the Short and Long trajectory, starting from a fixed reference state. If the speed is faster in the Short compared to the Long condition, we expect to have *t_long_* > *t_short_* for any arbitrary distance. Moreover, if the speed exactly scales with the average interval in each condition, we expect a linear relationship between *t_long_* and *t_short_*, with a slope equal to the predicted scaling factor, 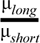 (Figure 2E).

When we applied this analysis to the neural data, we found that the empirical scaling factor accurately matched the predicted value (95% CI contained the predicted scaling factor: [1.39 1.58] for monkey G, [1.50 1.64] for monkey H; Figure 2F), both at the level of individual sessions and across animals (Figure S7). As a control, we generated a null distribution for the scaling factor after randomly reassigning the condition type (Short vs Long) for each neuron. In that case, we expected and observed no significant speed differences (95% CI for scaling factor: [0.78 1.26] for monkey G, [0.88 1.14] for monkey H; Figure 2F**, inset**). Note that because speed differences across conditions emerged ∼300 ms after Ready (Figure S4), we focused our analysis on the second half of the measurement epoch. However, results were robust when the analysis was extended over the entire measurement epoch (Figure S5). Together, these results provide compelling evidence that the speed of neural population dynamics within DMFC is adjusted according to the mean expected interval within each condition and reveal a precise mathematical relationship between temporal scaling and experienced temporal statistics in accordance with MPTS.

### Predictive temporal scaling is not explained by motor preparation

So far, we proposed that temporal scaling in the measurement epoch reflects a predictive process encoding the mean *t_s_*. However, since the RSG task requires *t_p_* to match *t_s_*, one plausible alternative explanation is that scaling is related to the average *t_p_*, and not the average *t_s_*. That is, scaling could reflect the fact that the animal is preparing to produce a short or a long interval (on average), depending on the condition. Indeed, many studies have found that when animals prepare a delayed motor response, the speed of dynamics during the motor preparation period scales with the instructed delay (*47, 48, 51, 55, 56, 65, 67–69*). It is therefore possible that the speed modulations in the measurement epoch are part of a global scaling that spans both Ready-Set and Set-Go epochs and is directly associated with the final motor preparation.

To test this alternative explanation, we analyzed neural recordings in DMFC in a variant of RSG, which we refer to as RSGgains, in which the *t_s_* and *t_p_* distributions were dissociated (*29*). RSGgains had the same basic skeleton as RSG: animals had to measure *t_s_* between Ready and Set, and produce *t_p_* between Set and Go. However, the key aspect that differentiated RSGgains from RSG was that the two conditions in which the animals performed the task had the same *t_s_* distribution but different *t_p_* distributions. Specifically, in one condition, *t_p_* had to match *t_s_* (same as RSG), while in a second condition, *t_p_* had to be equal to the measured *t_s_* multiplied by 1.5 (gain=1 or 1.5, respectively) (Figure 3A). Accordingly, RSGgains offered an ideal opportunity to verify that speed scaling during the measurement epoch was due to the distribution of *t_s_*, and not *t_p_*.

**Figure 3.**
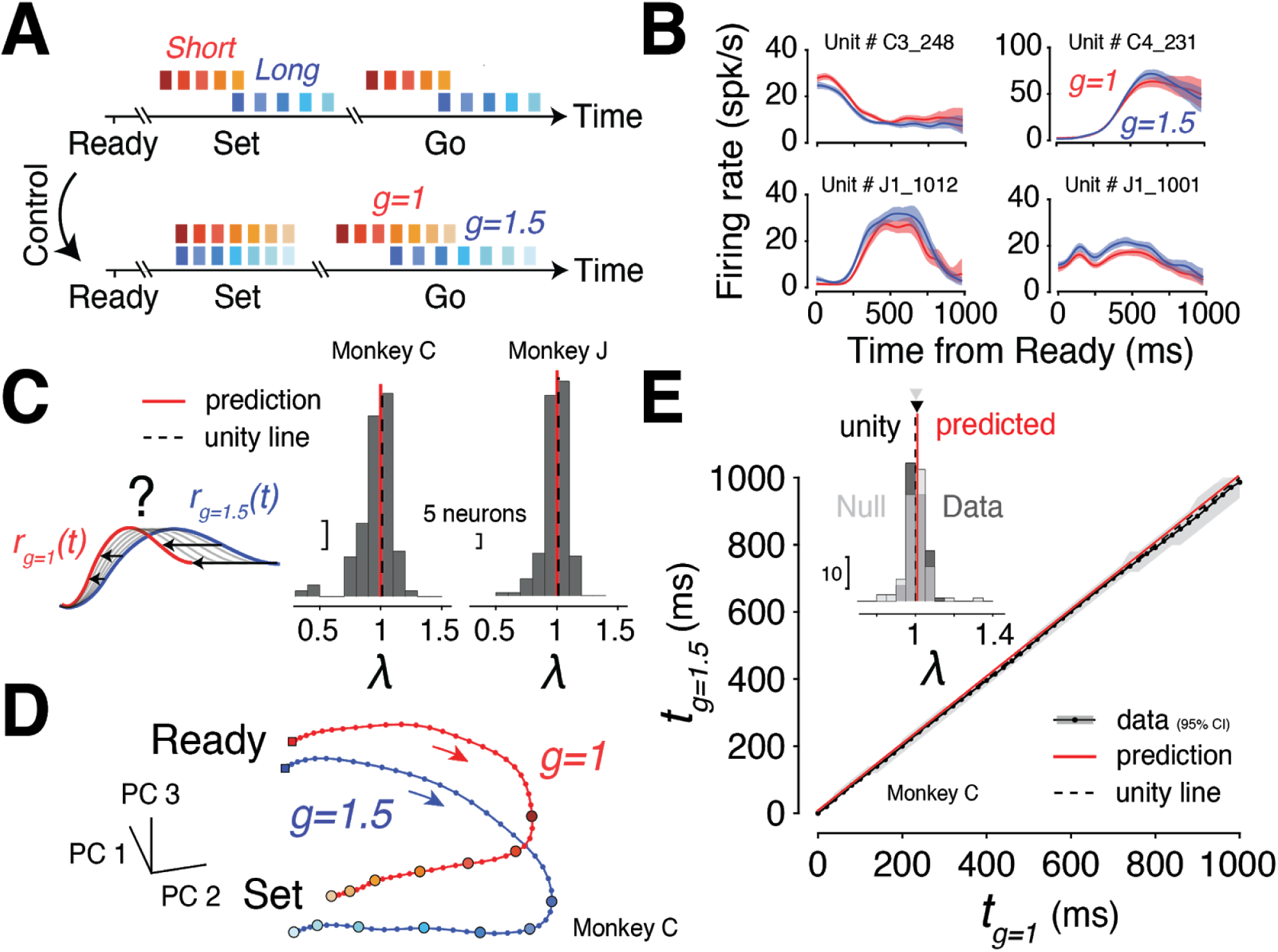
RSGgains experiment to rule out motor preparation as an alternative explanation for temporal scaling. **(A)** To verify that the neural speed during the measurement epoch of RSG reflected the underlying statistics of *t_s_*, and not that of *t_p_*, we used a control experiment (‘RSGgains’) to dissociate the *t_s_* and *t_p_* distributions. In contrast to regular RSG (top), in RSGgains (bottom) animals were exposed to two identical sample interval distributions but had to produce either 1 (gain=1, red) or 1.5 (gain=1.5, blue) times the measured interval (*29*). Similar to RSG, the gain condition was indicated to the animal by the color of the fixation point. **(B)** Firing rate of four example neurons during the measurement epoch of RSGgains (between Ready and Set), color coded by condition. Shaded areas denote 95% CI obtained by standard bootstrapping (N=100). Note the absence of temporal scaling compared to Figure 1B. **(C)** Scaling analysis at the single neuron level. We performed the same analysis as in Figure 2B on the RSGgains dataset (N=138 neurons for monkey C, N=201 for monkey J). When we fitted the firing rate of each neuron in the g=1 condition based on a scaled version of its firing rate in the g=1.5 condition, the distribution of scaling factors showed a single peak at one (black dashed line). This indicates that there were no systematic speed differences across conditions at the single neuron level. **(D)** Population dynamics in RSGgains. We plotted neural trajectories associated with both gain conditions in the space spanned by the top 3 PCs (∼90% of total variance explained). In contrast to RSG (Figure 2C), there is no apparent speed difference along the trajectories. **(E)** Empirical scaling factor. Similar to Figure 2F, we measured speed differences along the trajectories by computing the mapping between *t_g=1_* and *t_g=1.5_*, i.e., the time necessary to travel an arbitrary distance *d* on both trajectories. We estimated the empirical scaling factor by finding the slope of the mapping, and confirmed that both conditions evolved at the same speeds (data lies on the unity line, in agreement with the prediction; 95% CI for scaling factor: [0.87 1.12] for monkey C; [0.94 1.19] for monkey J). Inset: The regression slope of the mapping was not different from the null value (unity) obtained by shuffling conditions across neurons (unpaired *t*-test on bootstrapped distributions, *p*>0.05 for both monkeys).

We applied the same set of analyses to DMFC activity (138 neurons in monkey C, 201 in monkey J) in the RSGgains experiment. At the level of single neurons, activity during the measurement epoch did not exhibit any scaling effect between the two conditions (scaling factor across individual neurons, mean±sd, λ =0.96±0.14 for monkey C; λ =1.00±0.24 for monkey J; Figure 3B–C). At the population level, neural trajectories were separated between the two gain conditions (Figure 3D), similar to what we observed between the Short and Long conditions of the regular RSG task (Figure 2C). When we computed the speed along the trajectories, however, we found no speed differences between the two gain conditions (95% CI for scaling factor: [0.87 1.12] for monkey C; [0.94 1.19] for monkey J; Figure 3E, S8). This result is noteworthy, given that the only difference between RSG and RSGgains was whether or not the distribution of *t_s_* was the same in the two conditions. The presence of precise speed scaling in RSG and lack thereof in the RSGgains provides clear evidence that speed differences in the measurement epoch reflect the expected value of *t_s_*, and not *t_p_*.

### Behavioral adaptation to new temporal statistics

As a causal test for the MPTS hypothesis, we next sought to assess whether the speed of neural dynamics would change if the animals had to adapt to new temporal statistics. Indeed, one of the strong predictions of the theory is that predictive processing mechanisms should be adjusted flexibly to remain tuned to the current environmental statistics (*81–86*). We thus conceived another variant of RSG, which we refer to as RSGadapt, in which we first sampled *t_s_* from a fixed (‘pre’) distribution, and then covertly switched to another (‘post’) distribution (Figure 4A**, inset**). We chose the post-distribution to be a single interval at one extremum of the pre-distribution (i.e., shortest or longest interval). This choice was motivated by three factors. First, choosing the post-interval within the pre-distribution guaranteed that the transition to post could not be instantaneously detected. Second, having a single post-interval allowed us to track neural changes associated with the same sensory input during adaptation. Finally, we reasoned that this choice would enable us to leverage the bias in *t_p_* associated with the shortest and longest *t_s_* to track adaptation behaviorally. Specifically, we predicted that adaptation to a single *t_s_* at either end of the pre-distribution would be accompanied by a gradual removal of the bias associated with the reproduction of that *t_s_*. We also performed preliminary behavioral experiments in which the post-distribution included more than one interval (data not shown) but did not pursue that design as the behavioral adaptations were too slow and did not accommodate an analysis of neural changes within single electrophysiology sessions.

**Figure 4.**
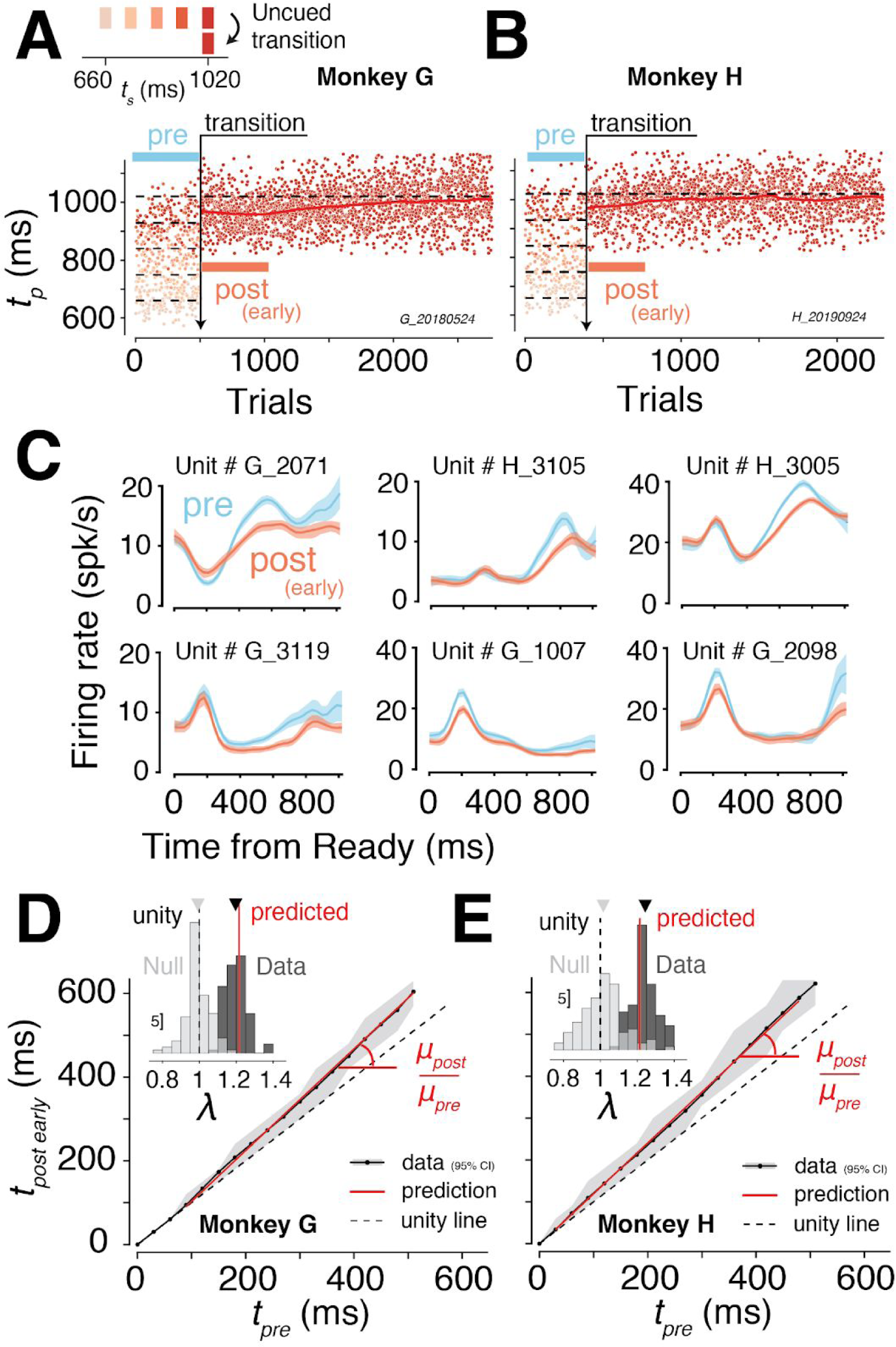
Behavioral and neural adaptation to changes in temporal statistics. **(A–B)** To assess how behavioral and neural responses would be affected by a covert change in temporal statistics, we challenged animals with a variant of RSG (called RSGadapt). In RSGadapt (top), we first sampled *t_s_* from a ‘pre’ distribution (660–1020 ms), and then covertly switched (in this case) to the longest interval of the pre-distribution (‘post’). The plot shows the time series of *t_p_* produced by the animal pre and post-switch. Each dot represents one trial, and *t_p_* are shown in chronological order, color-coded according to the associated *t_s_*. After the switch (vertical black arrow), *t_p_* gradually adjusted over thousands of trials until the initial bias toward the mean of the pre-distribution disappeared. The red line shows the running mean of *t_p_* post-switch (window: 300 trials). **(C)** Following the switch in interval statistics, we hypothesized that neural activity would adjust to reflect the new average interval. Because the mean of the post-distribution (μ*_post_* = 1020 ms) was longer than the mean of the pre distribution (μ*_pre_* = 840 ms), the MPTS hypothesis predicted that the speed should slow down by a scaling factor of λ = μ*_post_*/μ*_pre_* ≃ 1.21. The plot shows the firing rate of six example neurons pre-switch (blue) and early post-switch (dark pink). The number of trials pre and early post was matched (∼400 trials immediately before and after the switch). Changes in neural responses were qualitatively in line with our prediction: many of the neurons appeared to stretch their activity profiles post compared to pre-switch. **(D–E)** Empirical scaling factor early post-switch. Similar to Figure 2F, we measured speed differences between pre and early post activity by computing the mapping between *t_pre_* and *t_post early_*, i.e., the time necessary to travel an arbitrary distance *d* on both trajectories. The resulting mapping diverged from the unity line with a slope close to the predicted value μ*_post_*/μ*_pre_* (95% CI for scaling factor: [1.09 1.28] for monkey G, [1.11 1.38] for monkey H). Inset: distribution of scaling factors of bootstrapped data (black) and the corresponding null distribution (grey) obtained after randomly shuffling the condition (pre vs post) for each neuron.

We performed this manipulation in the same two monkeys used in the first experiment. Under the pre-distribution (*t_s_* between 660–1020 ms, mean: 840 ms), animals showed the characteristic biases toward the mean (Figure 4A–B). When we switched to the longest interval (*t_s_* = 1020 ms), animals’ responses immediately after the switch were indistinguishable from pre-switch (Wilcoxon rank sum test, *Z*=0.99*, p*=0.32 for monkey G, *Z*=-1.22*, p*=0.22 for monkey H), but then gradually adjusted over thousands of trials until responses were no longer biased. This was quantitatively confirmed by comparing the distribution of produced intervals early and late post-switch (two-sided unpaired t-test on *t_p_* distributions of the first and last 400 trials after the switch, *t*(1012)=-9.31, *p*<10^-10^ for monkey G, *t*(778)=-5.92, *p*<10^-8^ for monkey H). This adaptation was robust across animals and across experiments using either the shortest or the longest *t_s_* as the post-interval (Figure S9). One potential concern with using a single interval is that animals may choose to ignore the measurement epoch altogether and produce the desired *t_p_* from memory. We performed additional adaptation experiments that included catch trials to verify that animals continued to measure the single interval in the post condition (Figure S9).

### Neural adaptation to new temporal statistics

To examine the gradual changes associated with this adaptation at the neural level, we recorded DMFC activity (92 neurons in monkey G, 50 in monkey H) during a single session where animals experienced a change in interval statistics. Tracking changes of neural activity during learning is challenging because the recordings need to be 1) stable throughout the learning process, and 2) high-yield so that we can estimate neural states based on small numbers of trials during learning. Therefore, as a prerequisite, we first developed a custom recording approach that ensured the required stability and high-yield (see Methods and Figure S10).

To examine the neural correlates of learning, we first asked if the new temporal statistics had any consistent effect on the baseline firing rate of individual neurons. To do so, we computed the average firing rate of each neuron over the measurement epoch using ∼400 trials, both immediately before and after the switch. For the majority of neurons, mean firing rates had changed as a result of adaptation (72% (66/92) for monkey G, 58% (29/50) for monkey H, Figure 4C). However, similar to the two-context experiment, the direction of change was not systematic across neurons: on average, neurons were equally likely to increase or decrease their baseline firing rate between pre and post (two-sided paired t-test, *t*(91)=0.21, *p*=0.83 for monkey G, *t*(59)=-2.56, *p*=0.01 for monkey H; Figure S11). This result indicated that adaptation impacted firing rates, but the change did not involve systematic increase or decrease of the firing rates.

Next, we sought to test whether the activity changes were consistent with the temporal scaling of neural responses in proportion to the new mean interval (MPTS hypothesis). Given the mean of the post- and pre-distributions (μ*_post_* = 1020 ms and μ*_pre_* = 840 ms), MPTS predicts that the speed should slow down by a factor of 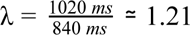. The slowing down of dynamics was already evident at the level of single neurons (Figure 4C), many of which appeared to stretch in time after the switch. To test the prediction quantitatively, we estimated speed differences at the population level between pre and post. Using the same analysis used in the two-context experiment, we found that the speed was adjusted to its predicted value robustly across the two monkeys (95% CI for scaling factor: [1.09 1.28] for monkey G, [1.11 1.38] for monkey H, Figure 4D–E). The data also rejected the alternative hypotheses according to which the speed might reflect either the shortest or the longest interval of the distribution. Indeed, the latter predicts that the speed should not have changed between pre and post 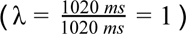 while the former predicts a scaling factor of 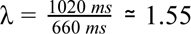. Both values were not included in the 95% confidence interval of the empirical scaling factor. Neural changes were thus fully consistent with animals adjusting their temporal expectations to the new distribution mean. This finding strengthens the validity of MPTS as it establishes a causal link between the interval mean and the speed of neural dynamics.

### Neural changes precede behavioral changes during adaptation to new interval statistics

At this point, we have established a tight mathematical relationship between changes in temporal statistics and neural adaptation: when the interval distribution changes, the speed of dynamics in DMFC is adjusted to quantitatively reflect the new mean interval. One important remaining question is whether the neural changes are caused by the new interval statistics or whether they reflect changes in the behavior, for example, through reafference (Figure 5A). If behavioral adaptation were to precede speed adjustments, we would not be able to reject the possibility that the neural changes in DMFC are a consequence of behavioral adaptation. In contrast, if speed adjustments were to precede behavioral adaptation, we would be able to conclude that neural changes are caused by the introduction of the new temporal statistics. To distinguish between these two possibilities, we examined the timescale of speed adjustments in DMFC to that of behavioral adaptation.

**Figure 5.**
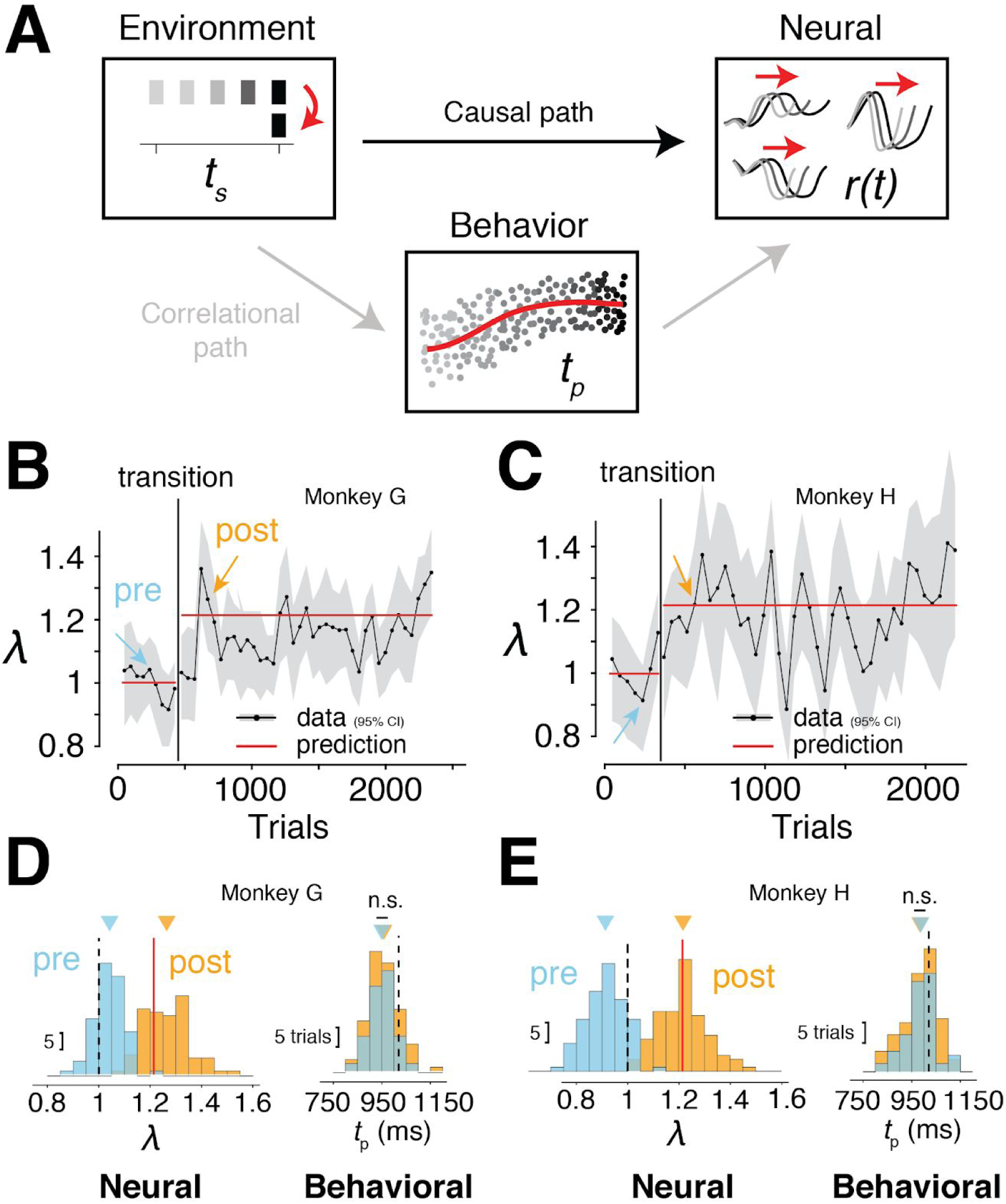
Timescale of neural and behavioral adaptation. **(A)** In the adaptation experiment, we found both behavioral and neural changes. One possibility (correlational path, in grey) is that neural changes were the indirect result of behavioral changes. For instance, changes in animals’ responses during adaptation to the new temporal statistics could affect neural activity through reafference and accumulate over trials to lead to persistent neural changes. Alternatively (causal path, in black), changes in statistics may have directly caused the observed neural changes, independent of behavioral adaptation. To dissociate between these two possibilities, we compared the timescale of neural and behavioral changes. Neural changes preceding behavioral changes would be consistent with the causal path, while behavioral changes preceding or co-occurring with neural changes would favor the correlational path. **(B–C)** To assess the timescale of neural adaptation following the switch in interval statistics, we examined how the empirical scaling factor changed in a 100-trial window (50% overlap) running throughout the session. To perform this analysis, we first computed a reference trajectory (*traj_ref_*) obtained by combining all trials (pre- and post-switch). We then computed the neural trajectory associated with each 100-trial block (*traj_block_*). To estimate the scaling factor associated with a particular block, we plotted the mapping between *t_block_* and *t_ref_*, corresponding to the time needed to travel an equal distance *d* along *traj_block_* and *traj_ref_*, respectively. We calculated the slope of this mapping (obtained via linear regression) for each block and normalized it by the average slope across all pre-switch blocks. Pre-switch slopes were thus expected to lie around unity, and subsequently reach their predicted value (λ = μ*_post_*/μ*_pre_*; red line). The data were in agreement with this prediction and showed that the speed rapidly adjusted to its predicted value. Shaded areas denote 95% CI obtained from bootstrapping. **(D–E)** A direct comparison of scaling factors (left) and produced intervals (right) pre-switch (blue arrow in panel B and C) versus post-switch (orange arrow) show that neural changes preceded behavioral changes. Roughly 200 trials following the switch, the neural speed had already adjusted (95% CI for scaling factor in the trial window, [1.13 1.44] for monkey G, [1.04 1.46] for monkey H), while animals’ behavior was virtually unchanged compared to pre switch (two-sided unpaired t-test on tp distributions, t(167)=0.97, p=0.33 for monkey G, t(162)=-0.38, p=0.71 for monkey H).

Our initial analyses already provided evidence that neural activity changed rapidly after the switch (i.e., ∼400 trials after the switch, Figure 4C). To quantify the timescale of neural changes rigorously, we computed how the scaling factor changed throughout adaptation using a 100-trial window running over the entire session (Methods). With this finer-grained analysis, we were able to confirm that ∼200 trials post-switch, the population speed was significantly different from pre-switch and not significantly different from its final predicted value (95% CI for scaling factor, [1.13 1.44] for monkey G, [1.04 1.46] for monkey H; Figure 5B–E). By comparison, animals’ behavior in the same post-switch trial window was statistically indistinguishable from pre-switch (two-sided unpaired t-test on *t_p_* distributions, *t*(167)=0.97, *p*=0.33 for monkey G, *t*(162)=-0.38, *p*=0.71 for monkey H, Figure 5D–E). This observation rules out the possibility that neural changes were purely driven by behavioral changes, and further supports the notion that changes in the neural speed reflected the dynamic adjustments of animals’ temporal expectations.

## Discussion

Predictive processing is among the most influential theories in neurobiology. Critical to this theory is the notion that the nervous system represents statistical regularities in the environment in the form of an internal model (*2–4*). The internal model encodes the predictable component of sensory inputs such that only residual errors from a priori predictions undergo further processing (*20–22*). However, the degree to which this theory can be applied broadly to explain neural mechanisms is still an active area of research (*26*). Our work extends and strengthens the case for this theory in a few important directions. First, current accounts of predictive processing are typically limited to modulations of static firing rates and thus cannot be straightforwardly extended to scenarios in which the information processing involves dynamics (*18, 57–63, 85*). Our work identifies a signature of predictive processing directly based on adjustments of neural dynamics. Second, prior work on predictive processing has been largely limited to information processing in the context of ‘where’ and ‘what’ questions (*13, 14, 20, 25, 87–91*), and not ‘when’ questions. This is somewhat surprising given the well-known capacity of humans to predict the timing of events in the future (*92*). By virtue of working in the domain of time, our work highlights the relevance of predictive processing in the temporal domain. Third, evidence for predictive processing has been largely limited to early sensory areas (*11, 23, 24*). Our work serves as an example of how this theory may explain neural observations in higher-order brain areas. Finally, and perhaps most importantly, our work provides a precise mathematical formulation of how neural signals precisely reflect the mean of temporal statistics.

Our results reveal a signature of predictive processing through the adjustment of neural dynamics based on experienced temporal statistics. In particular, we hypothesized and found compelling evidence that when awaiting a future event that can occur at different times, neural responses speed up or slow down in accordance with the statistical mean of the expected times. Converging evidence from several experiments made a strong case for this interpretation. First, the distribution of temporal scaling factors across single neurons in DMFC revealed a sharp and distinct peak at the value predicted by our hypothesis. Second, modulations of the speed of dynamics estimated by an unbiased analysis of neural trajectories across the population of neurons in individual animals and experimental sessions quantitatively matched the predictions of our hypothesis. Third, a control experiment verified that the modulation of speed was specifically related to the anticipated distribution of sensory events and not the ensuing motor response. Fourth, a novel learning experiment revealed that the speed adjustments were causally linked to temporal statistics of the environment. Importantly, in all these experiments, data was not consistent with the alternative hypothesis that temporal expectations are encoded by overall changes in baseline firing rates.

Like most discoveries, ours raises more questions than it answers. Here, we highlight some of the most pressing ones. First and foremost, what does the lawful relationship between neural speed and mean interval imply as a computational algorithm? This relationship establishes a form of invariance in the neural space: it guarantees that when the brain measures elapsed time, the neural trajectory may reach a fixed desired state at the expected mean interval irrespective of the underlying distribution. Consistent with this idea, we found evidence in our data that neural population activity reaches an invariant state at the mean interval and that a linear readout could decode deviations around this state to measure elapsed time directly in terms of prediction errors (Figure S12). Invariances of this kind are important as they could be the manifestation of a latent coding space that confers flexibility and generalization (*93, 94*). In the context of our finding, the brain may leverage this invariant distance metric to make relational inferences that generalize across contexts. For example, this organization would cause an interval that is one standard deviation away from the mean to be mapped onto the same neural distance to the mean irrespective of the distribution. Under suitable assumptions about noise, this scaling effect may provide a natural explanation for the scalar variability in time interval judgments (*95*). Similarly, this invariance may allow a criterion for categorical judgments to readily generalize to new stimuli (*96*). In our own work, the rapid adjustments of the neural speed in the RSGadapt learning experiment may have benefited from this representational invariance to perform rapid directed explorations in the neural space (e.g. meta-learning) (*97–102*). Specifically, modulations of speed based on the mean interval may serve to map time distributions onto a fixed range of desired states (*82, 85, 86, 103–105*), such that systematic changes in the mean would result in systematic prediction errors away from the desired states which in turn can drive efficient learning through directed exploration (*15*). Building on these observations, future work could examine whether this invariance is present and how it may facilitate rapid learning in other domains such as speed-accuracy tradeoff (*106, 107*) or the temporal control of attention (*78, 108*).

Juxtaposing our finding relating temporal scaling to the mean of an interval distribution with previous work reporting temporal scaling during time interval discrimination (*55, 65, 109*) and production (*28, 29, 47, 48, 56, 69, 73, 75, 110, 111*) raises the question of whether these phenomena are computationally related. We propose that all these observations may be unified under the theory of predictive processing, and the only thing that differentiate between them is the nature of what is being predicted. In our work, animals had to estimate a time interval, and thus it is natural for the system to predict the expected value of the distribution. In time interval discrimination task, scaling may accommodate the measurement of test intervals relative to the criterion. Finally, in motor timing tasks, scaling may serve as a prediction for the time of reward, which might be essential for reward-based learning and incorporating delay discounting information into reward prediction errors (*112, 113*). Scaling during motor planning may also play a role in predicting the sensory consequences of actions, which is thought to be an integral part of motor control (*114*). Our findings thus point to a unifying functional explanation of the phenomenon of temporal scaling commonly observed in timing tasks across species and brain areas (*43, 48, 55, 65–69, 75*).

In our experiments, we only tested uniform time distributions. It seems plausible that the results would hold for other symmetric unimodal distributions such as Gaussian. In contrast, for multimodal distributions, the mean interval may no longer be a good predictor. For example, for a bimodal distribution with two distinct modes, the system might adjust the speed to predict the two modes sequentially (*77, 115*) or use different groups of neurons (or subspaces) to generate predictions in parallel (*116–118*). In addition to predicting the mean, future work could investigate how the nervous system represents higher order moments such as variance and skewness. Currently, there is no definitive answer for how neurons encode variance although recent findings have suggested a functional role for nonlinear representations (*9, 28*). It is also not known how the brain represents skew in a distribution even though humans can be exquisitely sensitive to it (*119*). For skewed distributions, the scaling may continue to reflect the mean or may instead represent another statistic such as the mode (*120*). Finally, it is important to extend our findings to more elaborate sensorimotor tasks that involve both spatial and temporal uncertainty such as interception tasks (*121–123*), intuitive physics tasks (*124, 125*), or more complex agent-based pursuit tasks (*126–128*). These extensions will be critical for developing a comprehensive theory of how the nervous system learns spatiotemporal error distributions (*129*) and use that information to update internal predictions.

Finally, our learning experiment revealed that the nervous system learns to predict external events long before behavior can react to them. Influential theoretical (*130*) and behavioral studies (*131, 132*) have highlighted the potential utility of learning predictions before actions. Specifically, it has been hypothesized that predictions, which rely on *forward* internal models, serve to teach *inverse* models that control actions. Our work sets the stage for a direct experimental test of this hypothesis and provides a powerful platform for studying how predictive processing contributes to learning during sensorimotor behavior.

## Methods

All experimental procedures conformed to the guidelines of the National Institutes of Health and were approved by the Committee of Animal Care at the Massachusetts Institute of Technology. Experiments involved four awake behaving rhesus macaques (Macaca mulatta; ID: G and H for RSG2-prior and RSGadapt; weight: ∼8 kg; age: 5 years old; ID: C and J for RSGgains). Animals were head-restrained and seated comfortably in a dark and quiet room and viewed stimuli on a 23-inch monitor (refresh rate: 60 Hz). Eye movements were registered by an infrared camera and sampled at 1kHz (Eyelink 1000, SR Research Ltd, Ontario, Canada). The MWorks software package (https://mworks-project.org) was used to present stimuli and to register eye position. Neurophysiology recordings were made by 1 to 3 24-channel laminar probes (V-probe, Plexon Inc., TX) through a bio-compatible cranial implant whose position was determined based on stereotaxic coordinates and structural MRI scan of the animals. Analyses of both behavioral and electrophysiological data were performed using custom MATLAB code (Mathworks, MA).

### Behavioral task

#### RSG trial structure

Monkeys performed a time interval reproduction task known as the ‘Ready-Set-Go’ (RSG) task (*28, 133*). Each trial began with animals maintaining their gaze on a central fixation point (white circle: diameter 0.5 deg; fixation window: radius 3.5 deg) presented on a black screen. Upon successful fixation, and after a random delay (uniform hazard; mean: 750 ms, min: 500 ms), a peripheral target (white circle: diameter 0.5 deg) was presented 10 degree left or right of the fixation point and stayed on throughout the trial. After another random delay (uniform hazard; mean: 500 ms, min: 250 ms), the Ready and Set cues (white annulus: outer diameter 2.2 deg; thickness: 0.1 deg; duration: 100 ms) were sequentially flashed around the fixation point. Following Set, the animal had to make a proactive saccade (self-initiated Go) toward the peripheral target so that the produced interval (*t_p_*, between Set and Go) matched the sample interval (*t_s_*, between Ready and Set). The trial was rewarded if the relative error, |*t_p_*-*t_s_*|/*t_s_*, was smaller than 0.2. If the trial was rewarded, the color of the target turned green, and the amount of juice delivered decreased linearly with the magnitude of the error. Otherwise, the color of the target turned red, and no juice was delivered. The trial was aborted if the animal broke fixation prematurely before Set, or did not acquire the target within 3*t_s_* after Set. After a fixed inter-trial interval, the fixation point was presented again to indicate the start of a new trial. To compensate for lower expected reward rate in the Long condition due to longer duration trials (i.e., longer *t_s_* values), we set the inter-trial intervals of the Short and Long conditions to 1220 ms and 500 ms, respectively.

#### RSG2-prior experiment

In RSG2-prior (Figure 1 **and** 2), the sample interval, *t_s_*, was sampled from one of two discrete uniform distributions, with 5 values each between 480–800 ms for Short, and 800–1200 ms for Long. The distributions alternated in short blocks of trials (min of 5 trials for G, 3 trials for H, plus a random sample from a geometric distribution with mean 3, capped at 25 trials for G, 20 trials for H), and was indicated to the animal by the color of the fixation point (red for Short, blue for Long). On half the trials, and only in this first experiment, monkeys were required to acquire the peripheral target (at Go) using a hand-held joystick (‘hand trials’) instead of a saccade (‘eye trials’). However, for consistency with the other experiments (RSGgains and RSGadapt) which did not include hand trials, we only analyzed eye trials in this study.

#### RSGgains experiment

To verify that neural dynamics during the measurement epoch of the RSG task did not depend on the intervals produced by the animals, we analyzed neural recordings of two other monkeys (monkey C and J) performing a variant of RSG (‘RSGgains’; Figure 3). Full details of the task and experimental setup can be found in (*29*). Briefly, in this control experiment, *t_s_* was always sampled from the same discrete uniform distribution (7 values between 500–1000 ms) but animals had to apply a multiplicative gain (gain=1 or 1.5) to the measured interval before reproducing it. That is, animals had to produce either 1 or 1.5 times *t_s_*, depending on the condition. Similar to the 2-prior task, the condition type alternated in blocks of trials, and was indicated by the color of the fixation point.

#### RSGadapt experiment

In RSGadapt (Figure 4, 5 **and** 6), *t_s_* was first sampled from a pre-distribution (5 values between 660–1020 ms), and then became equal to a single interval, either the shortest or the longest value of the pre-distribution. The switch was not cued and occurred unpredictably during the session (typically between 500 and 800 trials after the start of the session to guarantee that there were enough trials both before and after the switch for physiology and for behavioral adaptation to happen with a single session). As a control, on some behavioral sessions, we introduced catch trials meant to probe animals on a different interval post-switch and verify that they were still measuring time even when exposed to a single interval (Figure S9). The catch trials were rare (∼6%) and unpredictable. Prior to recordings, and during behavioral training, animals experienced only a few of these adaptation sessions (12 out of 132 sessions over 239 days for monkey G, 17 out of 121 sessions over 274 days for monkey H), which were also interleaved with a series of ‘wash-out’ sessions with no switch. This was done to prevent overtraining the animals on these switches.

### Electrophysiology

#### Recording procedure

In RSG2-prior and RSGadapt, we used 2 to 3 laminar V-probes to record neural activity acutely in the dorsomedial frontal cortex (DMFC), comprising the supplementary eye field (SEF), the presupplementary motor area (pre-SMA), and the dorsal portion of the supplementary motor area (SMA). Figure S2 shows the exact electrode penetration sites on an MRI reconstruction of each animal’s brain. In RSGgains, recordings were done using 1 to 2 laminar V-probes inserted in the same area within DMFC (*29*). Signals were amplified, bandpass filtered, sampled at 30 kHz, and saved using the CerePlex data acquisition system (Blackrock Microsystems, UT). Spikes from single- and multi-units were sorted offline using the Kilosort software suite (*134*).

#### Recording stability

In RSG2-prior and RSGgains, experimental conditions (Short vs Long, gain=1 vs 1.5) were interleaved over relatively short blocks of trials (<100 trials); recording stability in these experiments was therefore not a concern when comparing neural activity across conditions (*28, 29*). In RSGadapt, however, we had to track neural activity over the course of an entire recording session (∼2-3 hours). We imposed a 1h settling time between electrode insertion and the onset of recordings to minimize electrode drift during the session. Figure S10 shows one example raster plot demonstrating stable spiking activity across trials. Recording stability was further confirmed by a more detailed analysis of spike waveforms (see Figure S10 for raw waveform traces) based on the approach used in (*135*). For every cluster extracted from spike sorting, we considered each spike waveform as a time series summarized as a point in a 60-dimensional space and computed the Mahalanobis distance between each waveform and the average waveform (across all spikes). This resulted in a distribution of distances, which we fitted to a chi-squared distribution following (*135*). We calculated the likelihood of each spike belonging to this fitted distribution, and any spike whose likelihood was lower than a fixed threshold was considered as an outlier. The threshold was defined as the inverse of the total number of spikes for that cluster. This analysis provided us with a single number (percent of outlier spikes) for every cluster, and allowed us to assess recording stability. In the analyzed RSGadapt sessions, we found that only 1% for monkey G, and 0% for monkey H of identified clusters had a percent of outlier spikes greater than 10%.

As an additional safeguard against instabilities, we systematically discarded clusters whose firing rate (averaged over the entire measurement epoch of the task) dropped below 1 spike per second. This guaranteed that neurons which disappeared over the course of the session (or were coming from noise) were not included in the analyses. The number of clusters (single or multi-unit activity) respectively for the RSG2-prior and RSGadapt experiments was 741 and 92 for monkey G, and 617 and 50 for monkey H. For the RSGgains experiment, the number of single/multi-units was 138 for monkey C, 201 for monkey J.

### Behavioral analyses

For behavioral analysis, we used a probabilistic mixture model to identify and reject outlier trials. Specifically, we calculated the likelihood of each *t_p_* (corresponding to a given *t_s_*) coming from either a task-relevant Gaussian distribution, or from a lapse distribution which we modelled as a uniform distribution extending from the time of Set up to 3*t_s_*. Any trial which was more likely to come from the lapse distribution was considered as an outlier (<5% trials) and discarded before further analyses.

### Single neuron analyses

Firing rates were obtained by binning spiking data (20-ms bins for RSG2-prior and RSGgains, 30-ms bins for RSGadapt), and smoothing using a Gaussian kernel (standard deviation of 40 ms). In addition, bootstrapped firing rates were generated via resampling trials with replacement (100 repeats). The number of trials where the visual target randomly appeared left or right of the fixation point were always matched before averaging across trials.

#### Estimating the rate of change of neural activity

To verify that the rate of change of neural activity (i.e., ‘speed’) was faster in the Short compared to the Long condition (Figure 1D), we computed the absolute difference in firing rates between consecutive 20-ms time bins and averaged these differences over the entire measurement epoch. We verified that the resulting speed difference across conditions was not due to the longer duration of the measurement epoch in the Long condition. When we matched the measurement epoch for the Short and the Long condition (i.e., restricting the analysis between Ready and the longest *t_s_* of the Short condition), speed differences were qualitatively unchanged.

#### Scaling analysis at the single neuron level

To assess how much single neuron activity profiles ‘stretched’ or ‘compressed’ in time across conditions (Figure 2A), we devised an analysis to reconstruct patterns of activity in the Short condition based on a scaled version of the pattern of activity in the Long condition. If we call *r_short_*(*t*) and *r_long_* (*t*) the pattern of activity of one neuron in the Short and the Long condition, respectively, our analysis searched for the set of parameters that minimized the difference [(γ *r_long_* (λ*t*) + δ) − *r_short_*(*t*)]^2^, where λ was the scaling factor, and γ and δ allowed for gain and baseline modulations across conditions. Example fits as well as distributions of fitted parameters are shown in Figure S3 **and** S6.

To better understand the source of the bimodality in the distribution of scaling factors across neurons (Figure 2B), we sub-selected neurons which had the largest firing rate modulations during the measurement epoch. We sorted neurons by the amount of variance in their firing rate across time points in the measurement epoch and kept neurons which contributed more than 0.1% of the total variance (150/741 for monkey G, 164/617 for monkey H).

#### Gaussian mixture model

To quantify the bimodality of the distribution of scaling factors across the population of neurons, we fitted a simplified Gaussian mixture model to the distribution. To do so, we first estimated the probability density of the empirical distribution (Gaussian kernel density estimator, *ksdensity* in Matlab). We then ran an optimization procedure (*fminsearch*) to minimize the root-mean-squared-error (RMSE) between the estimated probability density and the sum of two Gaussian probability density functions (pdf), each parametrized by their mean, variance and a gain factor multiplying the entire pdf.

### Population-level analysis

With the exception of the neural trajectories shown in the space of principal components (Figure 2C, 3D), all population-level analyses were performed using all the neurons (i.e., no dimensionality reduction was applied).

#### Temporal mapping analysis

To measure speed differences between neural trajectories (e.g., Short versus Long), we relied on a new ‘temporal mapping’ method inspired from the KiNeT analysis originally introduced in (*29*). For every state on the Short trajectory, we first computed the time elapsed (*t_short_*) and the distance traveled (*d_short_*) from a fixed reference state on the trajectory defined at *t_ref_* (relative to Ready). *d_short_* was calculated as the sum of the Euclidean distance between consecutive states along the trajectory between *t_ref_* and *t_short_*. Next, we found the state on the Long trajectory whose distance (*d_long_*) from the reference state was closest to *d_short_*, and marked the corresponding time (*t_long_*). Finally, we plotted *t_long_* as a function of *t_short_*, which we refer to as the *temporal mapping,* and computed the slope of this mapping to estimate the scaling factor related to speed differences across conditions.

Mathematically, if we call 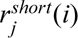 and 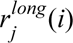 the firing rate of neuron *j* at time *i* respectively in the Short and Long condition, then for every *t_short_* > *t_ref_*, we calculated *t_long_* as follows:

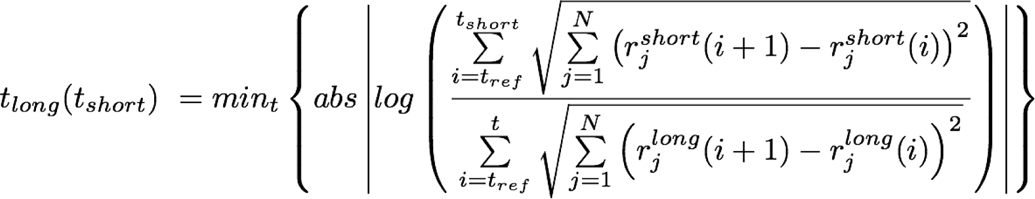

For RSG2-prior and RSGadapt (Figure 2F, 4D–E), the reference state was defined at 400 ms after Ready. This choice was motivated by the observation that, early after Ready, neural responses were largely dominated by a non-specific visual transient with equal speeds across conditions (Figure S4). We however ensured that our results were unaffected if the reference state was defined earlier in the measurement epoch (Figure S5). For RSGgains (Figure 3E, S8), where the speeds were hypothesized to be equal throughout the measurement epoch, the reference state was defined at Ready.

To compute the scaling factor associated with the two trajectories (i.e., slope of the temporal mapping), we proceeded in three steps. First, we identified the time point when the mapping started to diverge from the unity line (i.e., when *t_short_* becomes different from *t_long_*). We call this time point *t_onset_*. Second, we identified the time point when *t_long_* became equal to *max*(*t_short_*). We call this time point *t_offset_*. Finally, we used linear regression to estimate the scaling factor as the slope of the temporal mapping between *t_onset_* and *t_offset_*.

#### Running-window speed analysis

For the running-window speed analysis in the adaptation experiment (Figure 6B-C), we adjusted the analysis to ensure better statistical power. We first computed a global neural trajectory obtained by averaging population activity across all trials (pre and post-switch combined). We then computed the block-specific trajectory by averaging population activity in a 100-trial window (which we ran over the entire session; overlap between windows: 50%). The temporal mapping and the associated scaling factor (slope) were computed between the global trajectory and each block trajectories. Because the number of trials that went into each block trajectory was relatively small, individual temporal mappings were noisy. We therefore increased the number of points used in the regression analysis to compute the slope by placing the reference state at the time of Ready. Finally, we normalized the scaling factor in each block by the average scaling factor computed across all blocks during pre-switch. This normalization guaranteed that (1) the scaling factors post-switch could be directly tested against the predicted value μ*_post_*/μ*_pre_*, and (2) the potential bias introduced by including the early portion of the measurement epoch to compute the mapping slope was canceled out.

In one monkey (monkey H), the running-window speed analysis displayed large fluctuations toward the end of the session, i.e., after the neural changes had already converged to their predicted value. For this monkey and this analysis only, we relied on the waveform stability criterion to reject neurons which had more than 1% of outlier spikes (see Recording stability section above). This led to a decrease in the number of neurons included in the analysis (18 out 50) but was sufficient to resorb the large fluctuations.

#### Principal component analysis

PC trajectories (Figure 2C, 3D) were obtained by gathering smoothed firing rates in a 2D data matrix where each column corresponded to a neuron, each row corresponded to a given time point in the measurement epoch. To obtain a common set of principal components for different conditions (e.g., Short vs Long, or gain = 1 vs 1.5), we concatenated PSTHs for the different conditions along the time dimension. We then applied principal component analysis and projected the original data onto the top 3 PCs, which explained about 75% of total variance in RSG2-prior, and 85% in RSGgains.

## Acknowledgement

N.M. is supported by a MathWorks Engineering Fellowship and a Whitaker Health Sciences Fund Fellowship. H.S. is supported by a BBRF Young Investigator grant and the Center for Sensorimotor Neural Engineering. M.J. is supported by NIH (NIMH-MH122025), the Klingenstein Foundation, the Simons Foundation, the McKnight Foundation, and the McGovern Institute. We wish to thank Evan Remington for collecting the data for a control experiment, and Michal De-Medonsa for technical support. We also thank Seth Egger, Eli Pollock, Alexandra Ferguson and Sujaya Neupane for useful discussions.

## Supplementary Figures

**Figure S1.**
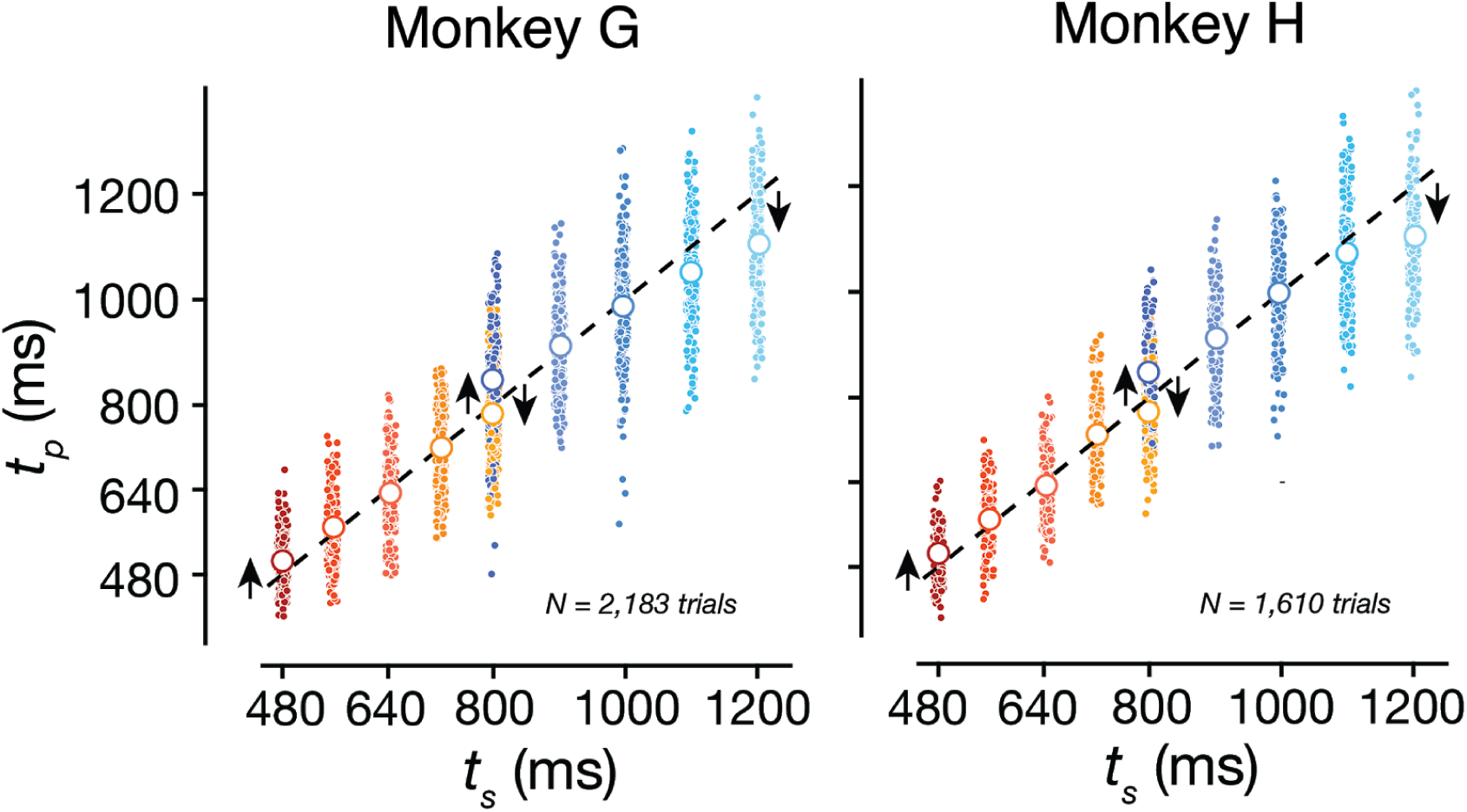
Monkey behavior in the RSG2-prior experiment. When exposed to two distributions of sample intervals (red for Short, blue for Long), animals systematically bias their responses toward the mean of each distribution (black arrows). Dots represent single trials; open circles show the average *t_p_* per each individual *t_s_*.

**Figure S2.**
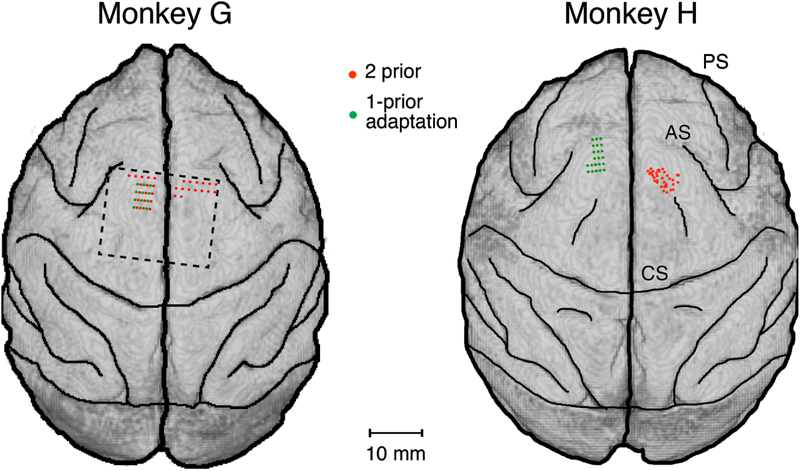
Recording sites for RSG2-prior and RSGadapt experiments. MRI surface reconstruction showing individual recording sites for both monkeys. Each dot represents one recording site; red for RSG2-prior experiment, green for RSGadapt. AS: arcuate sulcus; CS: central sulcus; PS: principal sulcus.

**Figure S3.**
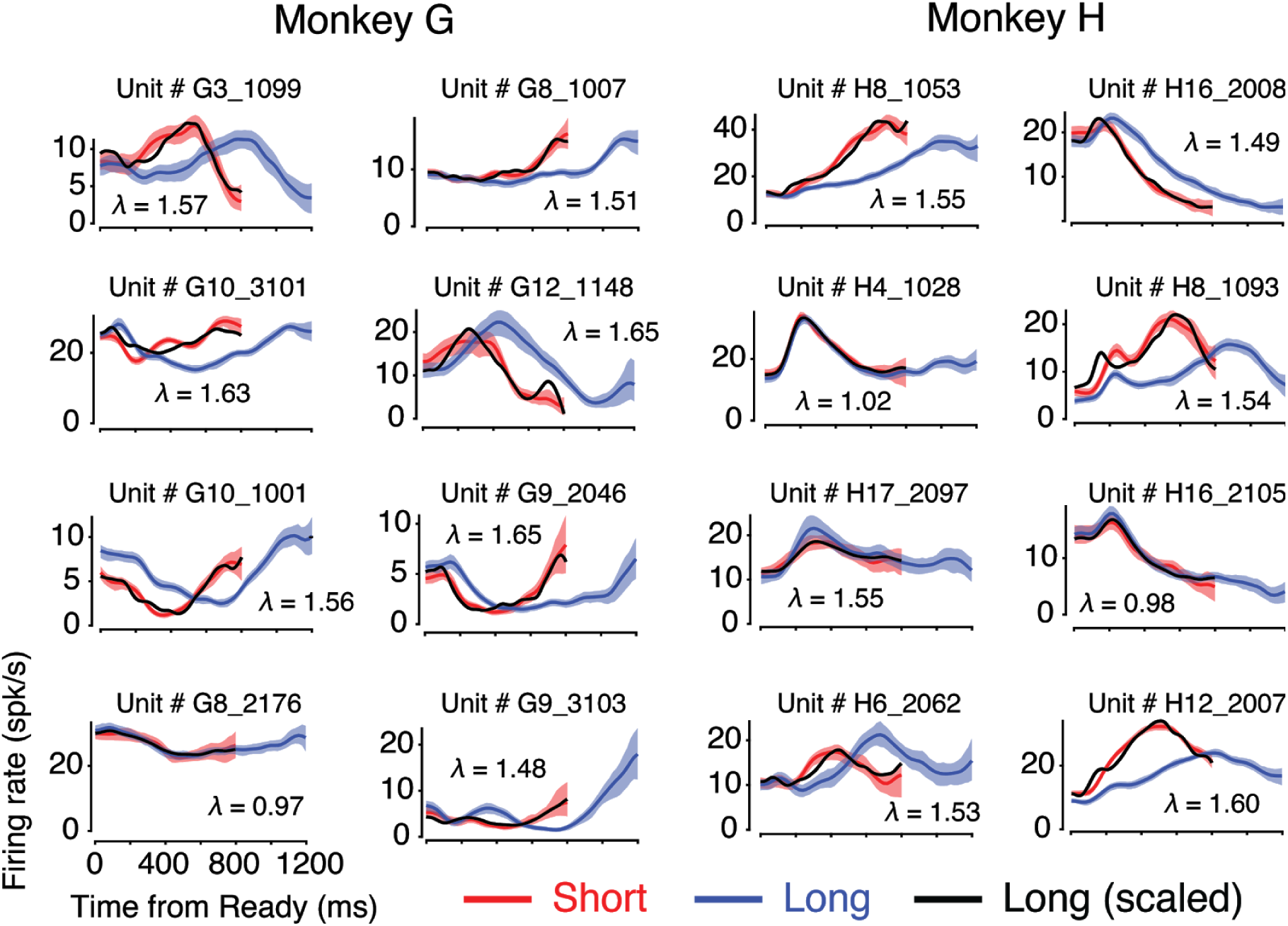
Single neuron activity during RSG2-prior measurement epoch. Red for Short, blue for Long, black for scaled version of Long fitted to Short (see Methods and Figure S6). Lambda is the fitted scaling factor. Shaded areas denote 95% CI obtained via standard bootstrapping.

**Figure S4.**
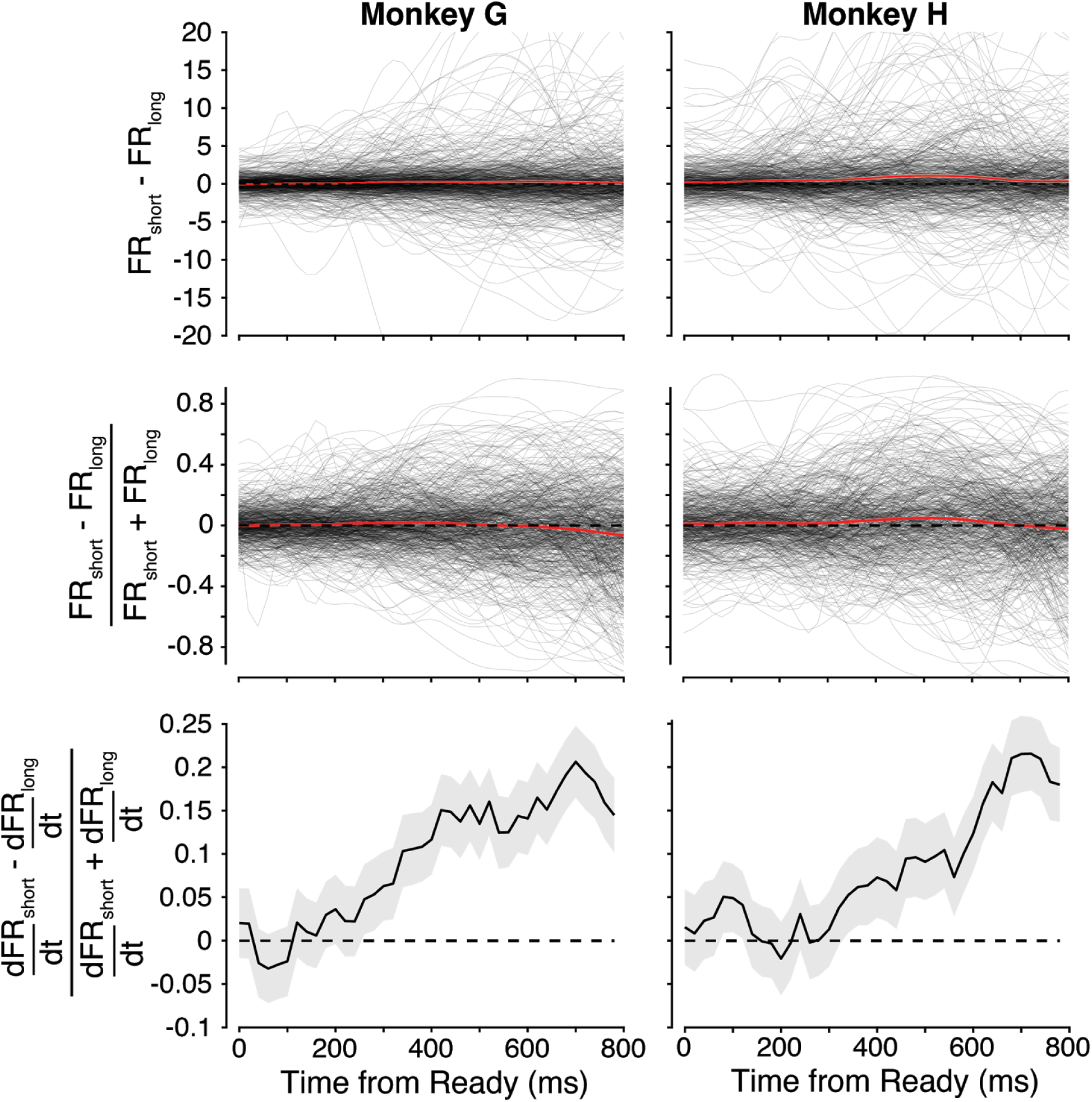
Differences in firing rate and speed across RSG2-prior conditions. Top: Difference in firing rates between the Short and Long condition throughout the measurement epoch of the task. Each black line represents one neuron; the red line represents the average difference across neurons. This difference fluctuated around zero and was unstructured across time points. Middle: Normalized differences in firing rates between the Short and Long condition throughout the measurement epoch of the task. Each black line represents one neuron; the red line represents the average difference across neurons. Similar to top, the normalized difference fluctuated around zero and was unstructured across time points. Bottom: Normalized differences in the rate of change of firing rates between the Short and Long condition. Black line shows average across neurons; shaded area denotes 95% CI.

**Figure S5.**
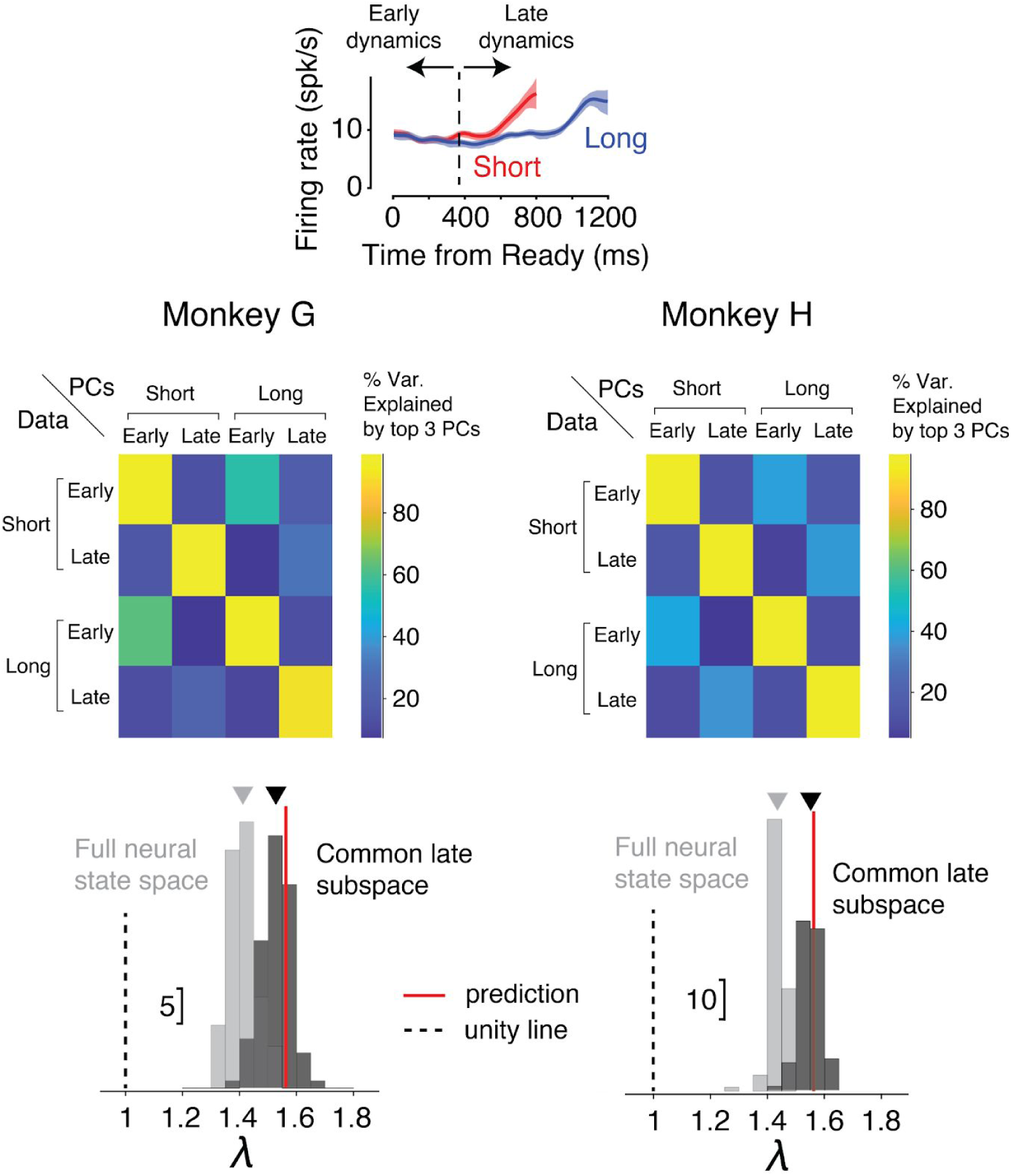
Early versus late dynamics during the measurement epoch of RSG2-prior. During the Ready-Set epoch of the task, neural responses had qualitatively different dynamics early (Ready – 400 ms) compared to late (400 ms – Set). Early dynamics tended to be non-condition specific, while late dynamics showed robust differences in speed across conditions (see upper panel for an example neuron; as well as Figure S4). To confirm this observation rigorously, we performed a cross-temporal principal component analysis (PCA) quantifying the amount of subspace overlap between the activity in the early vs late window of both Short and Long conditions. Results are summarized in a heatmap shown in the middle row: each cell (i, j) shows the percent of total variance explained when projecting the data of condition/window (i) onto the top 3 PCs computed from condition/window (j). Consistent with our qualitative observations, the overlap between early and late in each condition was smaller than the overlap across conditions for a given window. This result prompted us to define the reference state in our speed analysis (Figure 2F) at Ready+400ms to focus on the late part of the dynamics. Defining the reference state at Ready instead of Ready+400ms leads to a slight underestimation of the scaling factor (bottom row; grey distribution deviates from the predicted value 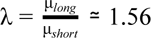; 95% CI [1.33 1.53] for monkey G, [1.39 1.48] for monkey H). This reflects the fact that early during the measurement epoch, dynamics is dominated by non-condition specific patterns of activity which bias the scaling factor towards unity. However, if we project the data onto the subspace associated with the late period of the measurement epoch (variance explained >85%), the resulting scaling factor matches the prediction even when the reference state is defined at Ready (black distribution overlaps with the predicted value; 95% CI [1.42 1.61] for monkey G, [1.48 1.61] for monkey H).

**Figure S6.**
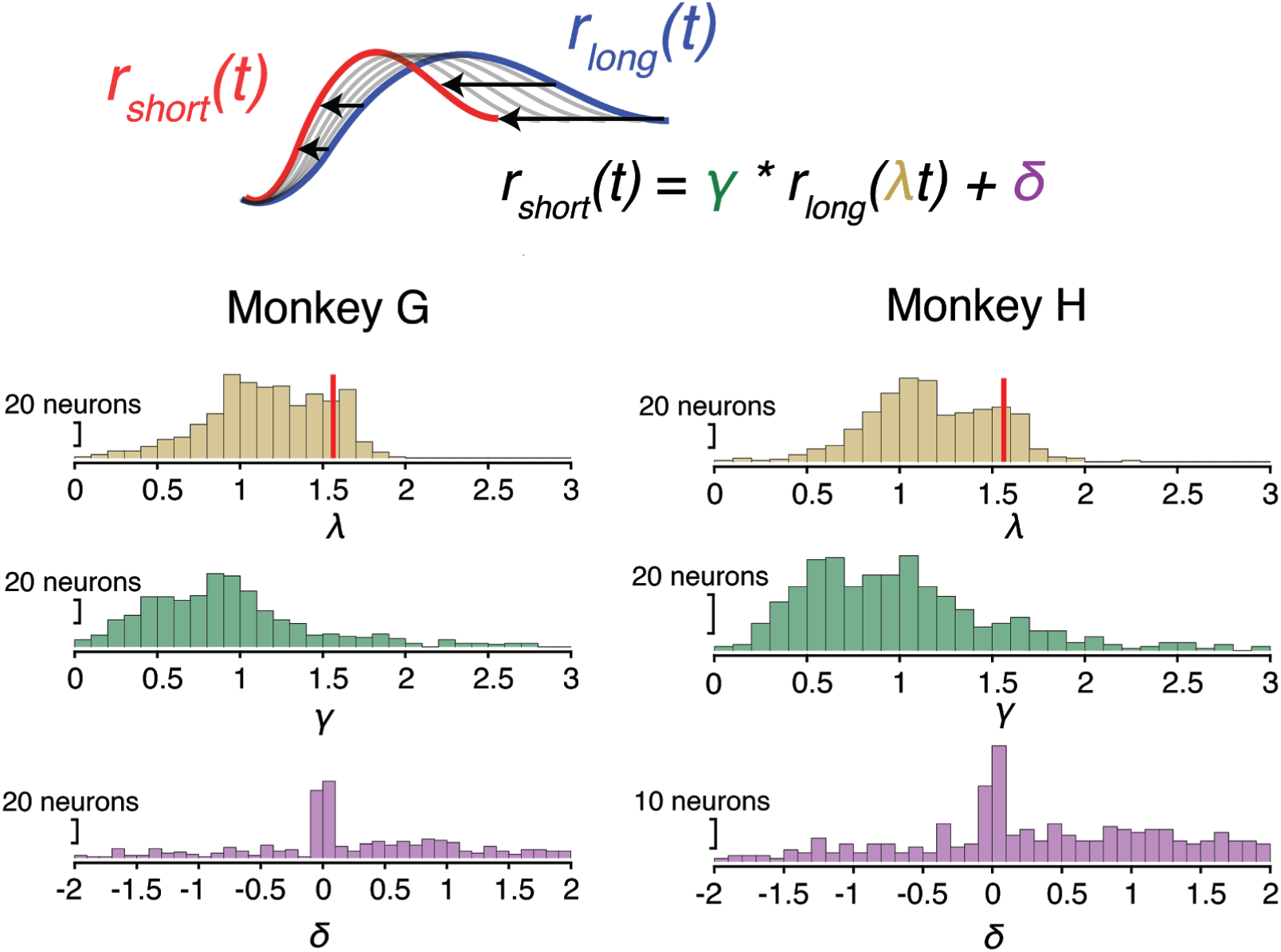
Parameter fits for temporal scaling in RSG2-prior. Top: each neuron’s activity profile in the Short condition (*r_short_*(*t*)) was fitted with a scaled version of its activity in the Long condition (*r_long_* (*t*)). The fitting procedure minimized the reconstruction error between *r_short_*(*t*) and γ *r_long_* (λ*t*) + δ, where λ is the scaling factor, and γ and δ allow for gain and baseline modulations across conditions. Bottom: distribution of fitted parameters across neurons in both monkeys. The red line on the λ distributions shows the predicted value 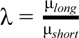, where μ*_short_* and μ*_long_* is the average time interval in the Short and Long condition, respectively.

**Figure S7.**
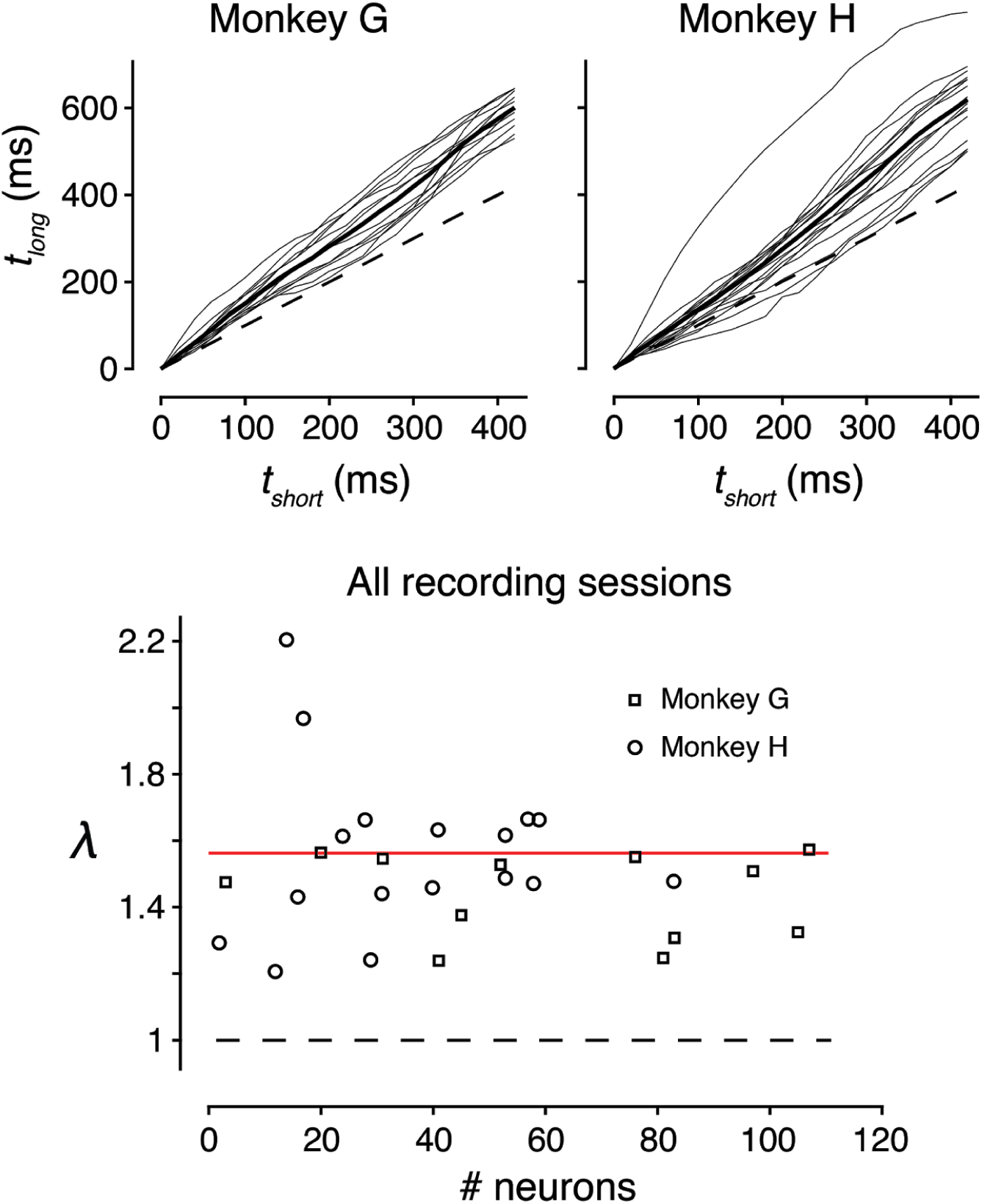
Temporal mapping across sessions in RSG2-prior. Top: each thin line represents one session (N=12 for monkey G; N=17 for monkey H), the thick line represents the average across sessions, the dashed line is the unity. Bottom: Scaling factor (mapping slope) for monkey G (squares) and monkey H (circles) as a function of recorded neurons in each session. Red line shows the predicted value 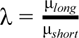.

**Figure S8.**
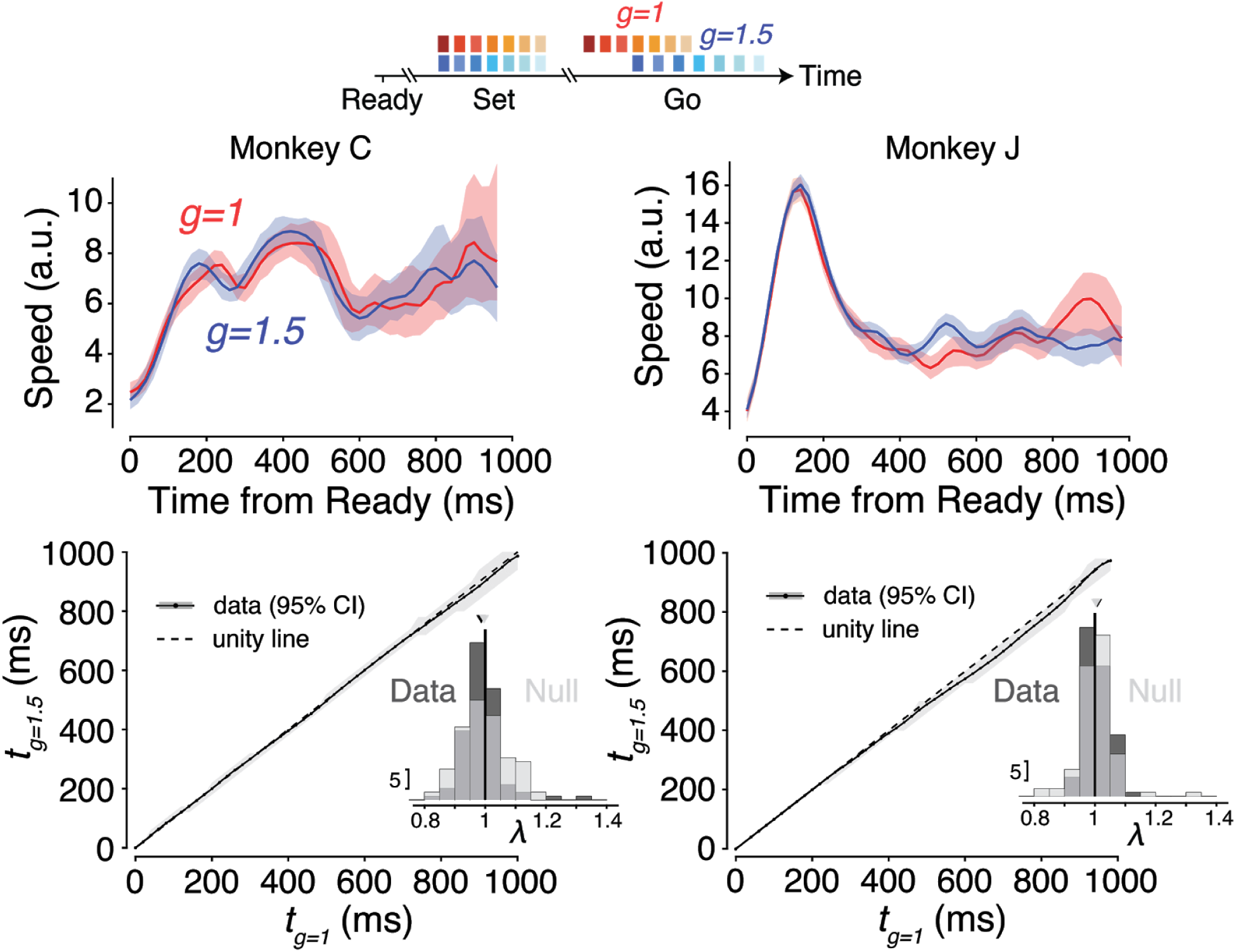
Temporal mapping and instantaneous speed in RSGgains. Top: absolute instantaneous speed was computed as the Euclidean distance between consecutive states separated by 20-ms bins. Throughout the measurement epoch, speeds did not diverge across conditions (red for g=1, blue for g=1.5). Bottom: temporal mappings for both monkeys lie along the unity (dashed) line, indicating no relative speed differences across conditions. Histograms show distributions of scaling factors for the bootstrapped data (dark grey) and a null distribution obtained by randomly shuffling conditions across neurons (light grey).

**Figure S9.**
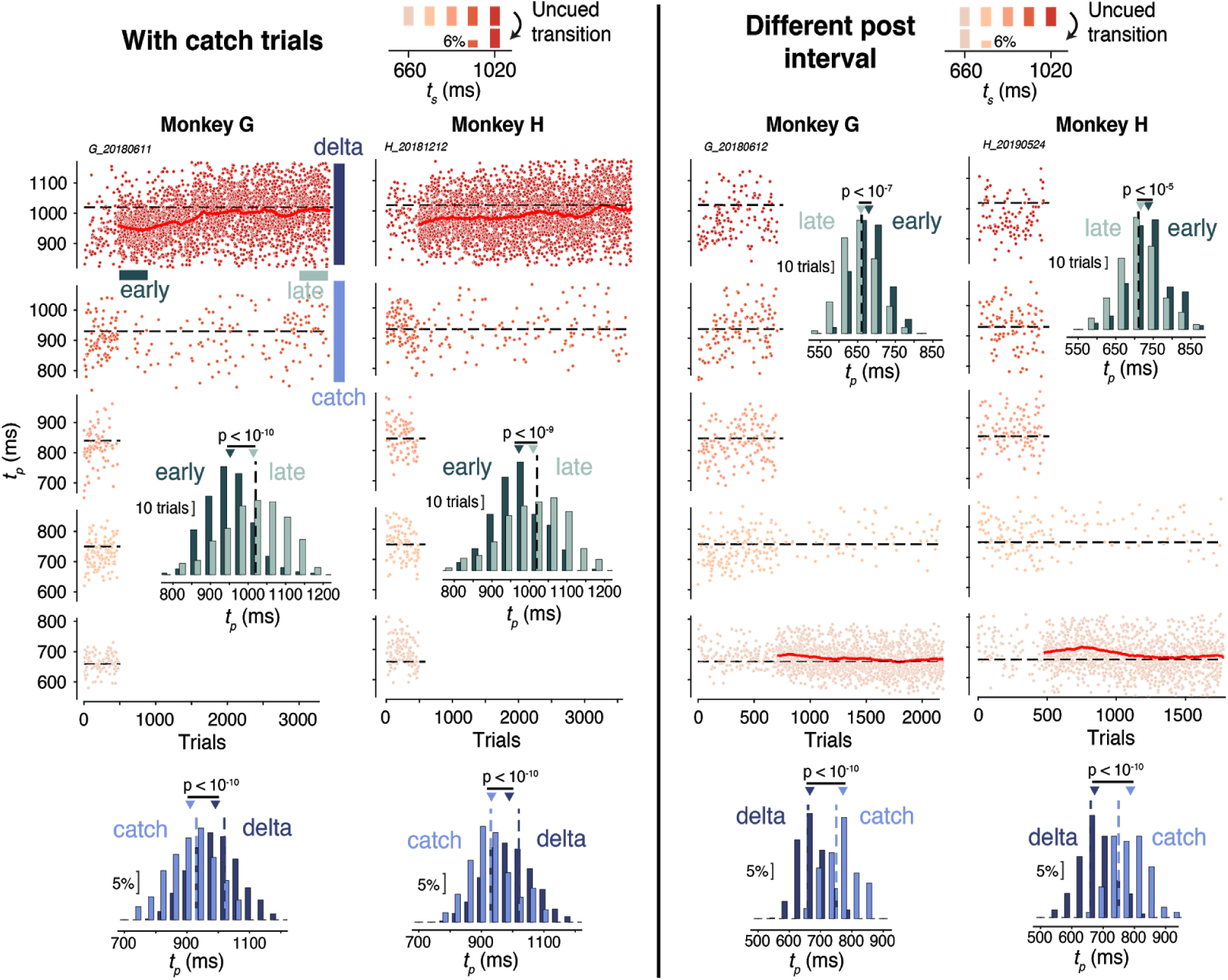
Monkey behavior on catch trials and different post intervals in RSGadapt. Left: to ensure animals were still measuring the sample time interval even when exposed to a single interval, we introduced rare (∼6%) ‘catch’ trials in which *t_s_* was different from the post (‘delta’) interval. In sessions where the delta interval was chosen to be the longest of the pre-distribution, the catch interval was chosen as the second longest interval of the pre-distribution. Bottom: *t_p_* distributions associated with delta (dark blue) and catch (light blue) trials were significantly different (two-sided unpaired-sample t-test, p<10^-10^ for both monkeys), indicating that animals continued to measure time while adapting to the new distribution (dark and light green distributions confirm the adaptation between early and late post-switch on the delta interval). Right: to test the robustness of behavioral adaptation, we ran sessions in which the post-interval was the shortest interval of the pre-distribution. Comparison of *t_p_* distributions early (dark green) and late (light green) post-switch confirmed that the animals successfully adapted (p<10^-5^ for both monkeys). Introduction of catch trials in these sessions further confirmed that animals continued to measure time post-switch.

**Figure S10.**
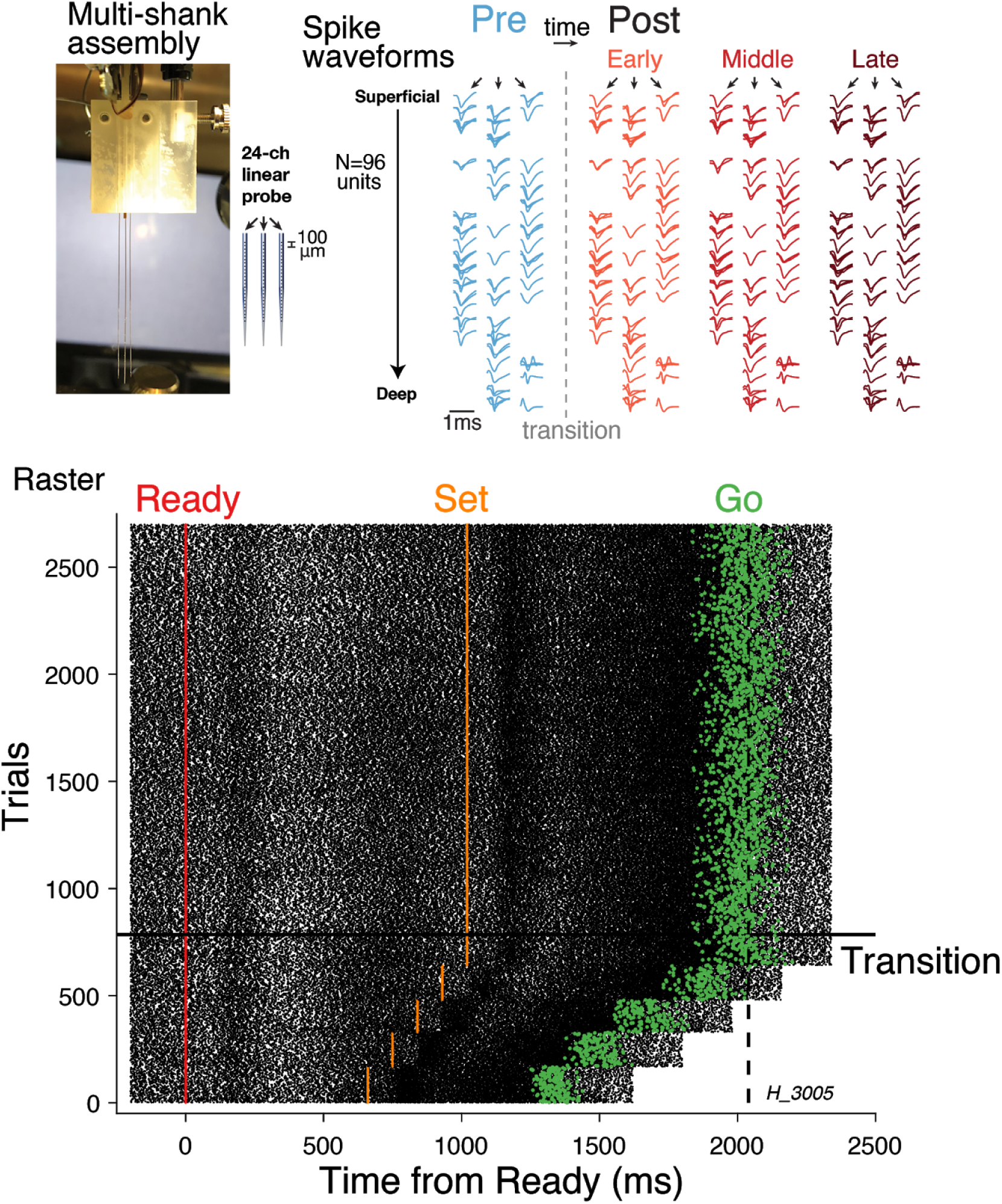
Recording stability and example raster plot during RSGadapt. Top: To verify the stability of our recordings, we plotted the waveforms at different stages of the recording session during the adaptation experiment (blue for pre-switch; light, intermediate, dark orange for early, middle, late post-switch). For visualization, waveforms are spatially arranged to match their recording sites on the three V-probes (superficial to deep from top to bottom). Bottom: one example neuron’s raster plot (see Figure 4C for the PSTH associated with that unit). Spikes are aligned to Ready (red); orange dots show Set, green dots show Go. For clarity, pre-switch trials have been sorted by *t_s_*, however in the experiment, *t_s_* was randomly sampled. Trials post-switch (above horizontal black line showing the transition) are shown in chronological order.

**Figure S11.**
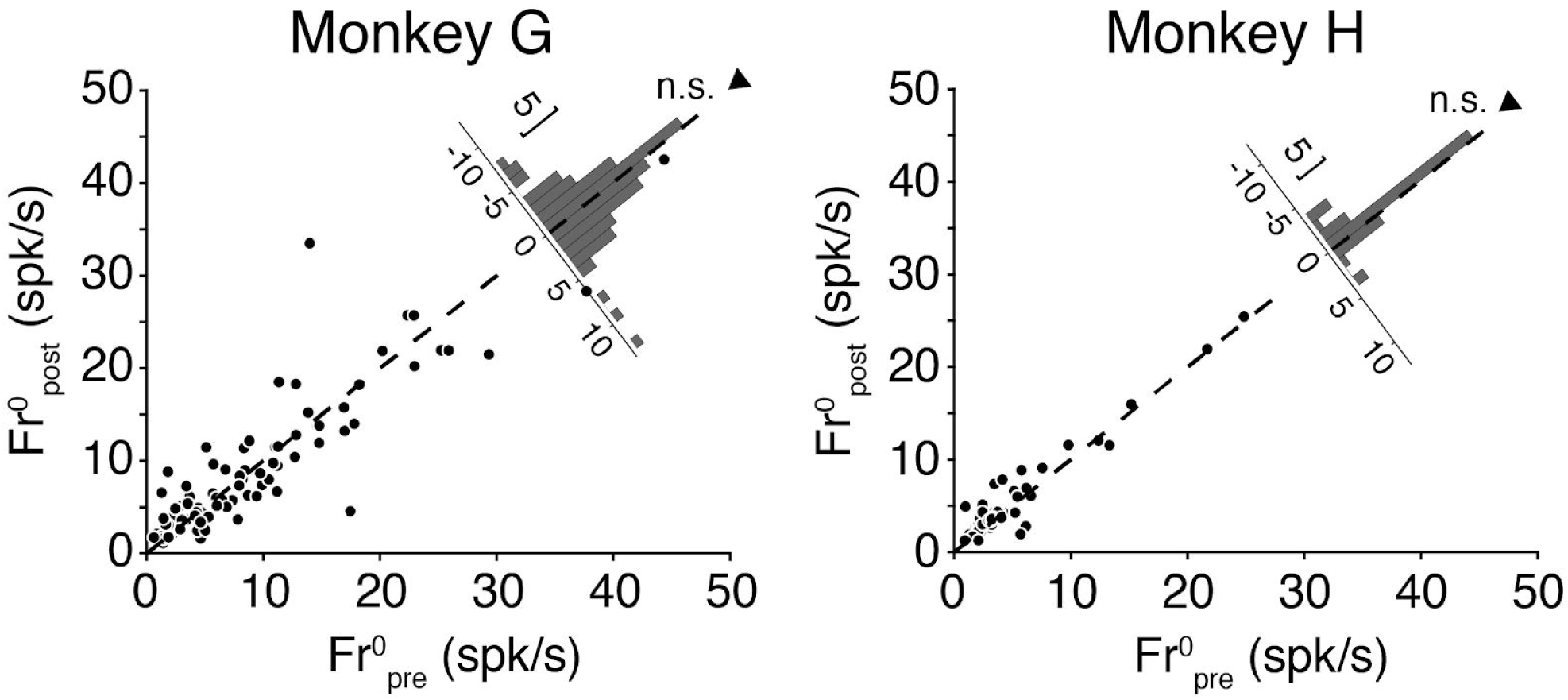
Differences in firing rate before and during adaptation in RSGadapt. We plotted the baseline activity of each neuron at the time of Ready (20-ms window following Ready) in the pre (abscissa) and early post (ordinate) condition. Pre and early post had the same number of trials (∼400 trials immediately before and after the switch). Each dot represents one neuron. The diagonal distribution shows that there were no systematic differences in firing rates across the population (*p*>0.01 for both monkeys).

**Figure S12.**
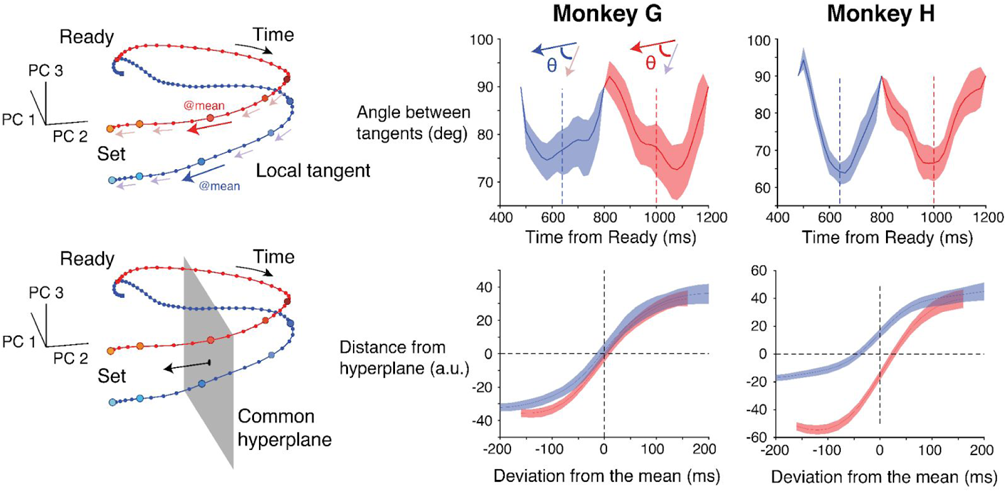
Invariance of neural states across conditions in RSG2-prior. Speed modulations across conditions based on the mean interval may allow the neural trajectory to reach an invariant state at the mean expected time of Set irrespective of the underlying time distribution. Deviations away from this state at the time of Set may in turn allow the system to measure intervals in terms of a prediction error. We performed two complementary analyses to test this idea. First, we asked whether neural dynamics near the state associated with the mean interval (i.e., mean state) was self-similar across conditions (top). To do so, we computed the local tangent at the mean state for each trajectory (i.e., mean tangent; large red and blue arrows in the left panel). We then calculated the angle between the mean tangent of one condition and the local tangents computed along the trajectory of the other condition (smaller pink and purple arrows). For any given state, the local tangent was computed by connecting the states immediately preceding and following that state (bin size: 20 ms). In both animals (middle and right columns), we found that the angle was minimum when the tangents were computed near the mean state, suggesting that local dynamics was most similar around the two mean states. Next, we asked whether we could define a hyperplane slicing the two trajectories near the mean state to linearly decode elapsed time in terms of prediction errors (bottom). Importantly, the hyperplane needed to be *common* to both trajectories if the system is to leverage a shared coding scheme to measure prediction errors irrespective of the time distribution. We chose the hyperplane (grey plane) which contains the average of the two mean states and whose normal vector (black vector) is the average of the two tangent vectors at the corresponding means. We then calculated the distance to the hyperplane for each neural state along both trajectories (by convention, the distance was positive when the state lay on the side of the hyperplane toward which the normal vector pointed). Finally, we plotted distance to the hyperplane as a function of elapsed time relative to the mean interval in each condition separately. For both monkeys, distance vs relative time had a monotonic profile, confirming that a common hyperplane can be used to linearly decode time intervals as prediction errors away from the mean.

